# *Shigella flexneri* evades LPS ubiquitylation through IpaH1.4-mediated degradation of RNF213

**DOI:** 10.1101/2024.09.24.614686

**Authors:** Katerina Naydenova, Keith B. Boyle, Claudio Pathe, Prathyush Pothukuchi, Ana Crespillo-Casado, Felix Scharte, Pierre-Mehdi Hammoudi, Elsje G. Otten, Felix Randow

**Author notes:** These authors contributed equally (Katerina Naydenova, Keith B. Boyle, Claudio Pathe).

## Abstract

The evolutionary arms race between pathogens and hosts has resulted in pathogens acquiring diverse adaptive countermeasures that antagonize host immunity. Ubiquitylation of lipopolysaccharide (LPS) on cytosol-invading bacteria by the E3 ligase RNF213 creates ‘eat-me’ signals for antibacterial autophagy but whether and how cytosol-adapted bacteria avoid LPS ubiquitylation remains poorly understood. Here we show that *Shigella flexneri*, a professional cytosol-dwelling enterobacterium, actively antagonizes LPS ubiquitylation through IpaH1.4, a secreted effector protein with ubiquitin E3 ligase activity. IpaH1.4 binds to the LPS E3 ubiquitin ligase RNF213, ubiquitylates it, and targets it for degradation by the proteasome, thus preventing LPS ubiquitylation. To understand how IpaH1.4 recognizes RNF213, we determined the structure of their complex using cryogenic electron microscopy. The specificity of the interaction is achieved via the leucine rich repeat of IpaH1.4, which binds the RING domain of RNF213 by hijacking the conserved RING interface required for binding of ubiquitin-charged E2 enzymes. Interestingly, IpaH1.4 also targets the E3 ligase LUBAC - required for the synthesis of M1-linked ubiquitin chains on cytosol-invading bacteria downstream of RNF213 – as well as multiple other E3 ligases involved in inflammation and immunity – through binding to the E2-interacting face of their RING domains. We conclude that IpaH1.4 has evolved to antagonize multiple anti-bacterial and pro-inflammatory host E3 ligases.

## Introduction

The intracellular lifestyle has evolved multiple times amongst pathogenic bacteria, enabling them to gain access to cellular nutrients and also to avoid potent extracellular immunity.^1^ However, most intracellular bacteria reside in membrane-delimited compartments with only a select few succeeding in colonizing the host cytosol.^2^ Access to cellular nutrients is arguably easier in the cytosol, suggesting that cytosolic anti-bacterial immunity is particularly potent as it successfully prevents colonization by most facultatively intracellular bacteria. To sense and curb bacterial infection in the cytosol, a variety of cell-autonomous immune mechanisms exist. These include inflammasome activation, the induction of cell death, and anti-bacterial autophagy. The lattermost relies on pairs of ‘eat-me’ signals associated with bacteria and cognate receptors that promote autophagy^3^, for example sphingomyelin on stressed but otherwise intact bacteria-containing vacuoles sensed by TECPR1^4^, glycans on broken bacteria-containing vacuoles sensed by galectin-8 / NDP52^4^, as well as ubiquitin conjugated to the bacterial surface and sensed by multiple cargo receptors including NDP52, TAX1BP1, Optineurin and SQSTM1.^5–9^

We recently discovered that the initial step of ubiquitin coat formation on cytosol-exposed *Salmonella enterica* serovar Typhimurium is catalyzed by the E3 ligase RNF213, which ubiquitylates bacterial lipopolysaccharide (LPS) in the first example of ubiquitylation targeting a non-proteinaceous substrate.^10^ Ubiquitylation of LPS results in the downstream recruitment of LUBAC, a multimeric E3 ligase that adds M1-linked ubiquitin chains to RNF213-initiated ubiquitin coats.^11,12^ In addition to Gram-negative bacteria, RNF213 restricts also pathogens that do not produce LPS, including Gram-positive bacteria, the apicomplexan parasite *Toxoplasma gondii* and certain viruses.^13–18^ Mutations in RNF213 cause Moyamoya disease, a rare cerebrovascular disorder caused by intimal thickening and occlusion of the terminal portion of the internal carotid artery.^19–21^

Given that some bacteria successfully colonize the host cytosol, we wondered how they escape LPS ubiquitylation. *Shigella flexneri* is an example of a cytosol-dwelling Gram-negative bacterium that has evolved a plethora of mechanisms to avoid clearance by cell autonomous immunity, including anti-bacterial autophagy, and to establish its replicative niche in the cytosol.^22–24^ These adaptations include the IpaH effector proteins, a family of bacterial E3 ubiquitin ligases, that are secreted into the cytosol of infected cells, where they engage the host ubiquitin-proteasome system.^25–28^ IpaH proteins comprise a short unstructured N-terminus, a leucine-rich repeat (LRR) mediating substrate recognition, and a C-terminal, so-called ‘novel E3 ligase’ (NEL) domain. It has been suggested that in the absence of substrate IpaH ligases are auto-inhibited by their LRR domain through blockade of the catalytic cysteine in the E3 domain.^29–32^ *S. flexneri* encodes up to twelve functional IpaH proteins, five of which are found on the *Shigella* virulence plasmid: *ipaH1.4*, *ipaH2.5*, *ipaH4.5*, *ipaH7.8*, *ipaH9.8*. Examples of IpaH ligases and their immuno-regulatory host substrates include IpaH9.8 and guanylate-binding proteins (GBPs)^33–35^, IpaH7.8 and gasdermin B / gasdermin D^36–38^, IpaH9.8 and NEMO^39^ as well as IpaH1.4 and LUBAC^40,41^. In all known cases, the IpaH ligase targets important anti-bacterial host proteins for proteasomal degradation through conjugation of polyubiquitin chains. IpaH proteins thus synergistically enable *Shigella* to retain actin-dependent motility, to block pyroptotic cell death and to reduce NF-κB activation, together dampening the anti-bacterial and pro-inflammatory response of their host cells.

In this study, we investigate whether *S. flexneri* antagonizes LPS ubiquitylation in the cytosol of infected cells. We find that *S. flexneri* does indeed not suffer from LPS ubiquitylation because RNF213, the E3 ligase catalyzing LPS ubiquitylation^10^, is bound, ubiquitylated, and targeted for proteasomal degradation by the bacterial effector protein IpaH1.4. Cryogenic electron microscopy revealed that IpaH1.4 binds to the RING domain of RNF213. Remarkably, *S. flexneri* deploys the same IpaH1.4 effector to also target a RING domain in LUBAC^40,41^ (known to produce M1-linked ubiquitin chains on bacteria downstream of RNF213^11^), and in several other RING-containing proteins, thus revealing how a single bacterial effector antagonizes multiple steps in a complex cascade of anti-bacterial host proteins.

## Results

### *Shigella flexneri* produces a trans-acting factor to avoid LPS ubiquitylation

We previously demonstrated that the facultatively cytosol-dwelling bacterium *Salmonella* Typhimurium undergoes RNF213-dependent LPS ubiquitylation.^10^ To test whether a professionally cytosol-invading bacterium is similarly recognized, we infected HeLa cells, a human epithelial cell line, with *Shigella flexneri.* We noticed that, unlike *Salmonella* Typhimurium, *Shigella flexneri* did not become ubiquitin-coated in the host cytosol (**Figure 1A**). To directly address LPS ubiquitylation, we immunoblotted lysates of bacteria isolated from cells at 4h post infection with an antibody specific for conjugated ubiquitin (FK2) (**Figure 1B**). Immunoblotting revealed an LPS-ubiquitin smear and a characteristic banding pattern in the positive control samples of wild type and *ΔrfaL Salmonella* Typhimurium, respectively, which are due to the synthesis of high and low molecular weight LPS in the two strains.^10^ In contrast, ubiquitylated LPS was not detected in lysates from cells infected with either wild type or *ΔrfaL Shigella flexneri*. In co-infection experiments, *Salmonella* was protected from ubiquitylation when present together with *Shigella* in the same host cell (**Figure 1C**). This observation suggests the existence of a secreted *Shigella* effector acting *in trans* to antagonize LPS ubiquitylation on *Salmonella*. We also noticed that lack of ubiquitylation of *S. flexneri* correlated with lack of recruitment of endogenous RNF213 to the bacteria (**Figure 1A, C)** and that recruitment of RNF213 to the surface of cytosolic *Salmonella* was abolished in cells co-infected with *Shigella*. We therefore conclude that the trans-acting factor produced by *Shigella* directly antagonizes RNF213 and is not merely removing ubiquitin from bacteria.

**Figure 1.**
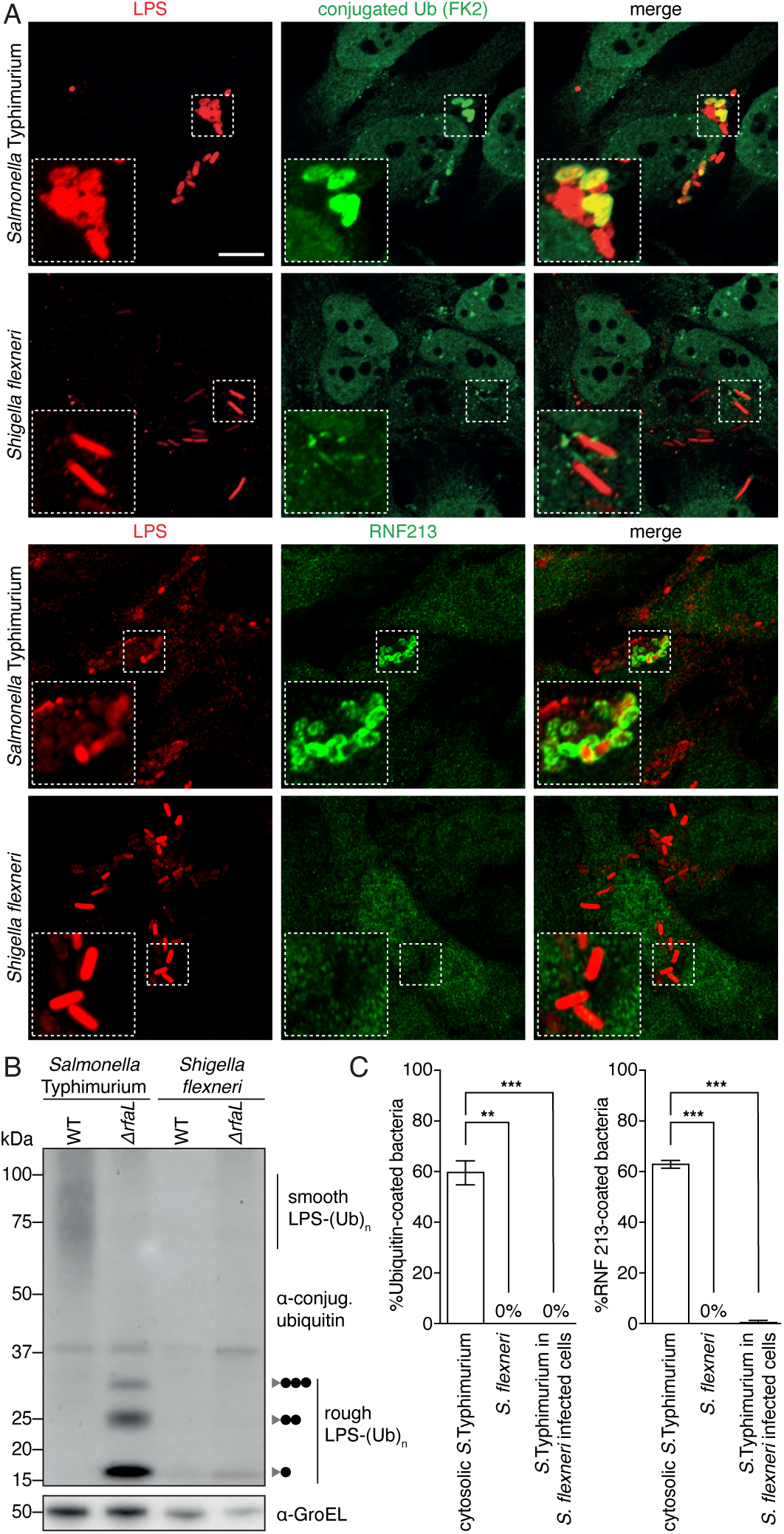
A *Shigella*-derived trans-acting factor antagonizes LPS ubiquitylation by RNF213. (A) Representative confocal micrographs of HeLa cells at 5 hours post infection with *Salmonella* Typhimurium (upper) or *Shigella flexneri* (lower), immunostained for LPS (red) and conjugated ubiquitin (FK2) (green), or LPS (red) and RNF213 (green) as indicated. Dashed squares, representative areas magnified 2.5× in the insets. Scale bar, 10 μm. (B) Immunoblot analysis of the indicated strains of S. Typhimurium and *S. flexneri* extracted from HeLa cells at 5h post infection. WT, wild type; grey triangle, *ΔrfaL* LPS; black circle, ubiquitin; GroEL, loading control for bacterial lysates. The LPS fractions were isolated by heat clearance of bacterial lysates and probed for conjugated ubiquitin with FK2 antibody. The loading control (GroEL) was probed for in non-heat cleared bacterial lysates. (C) Percentage of bacteria positive for conjugated ubiquitin (detected by FK2 antibody) (left panel) and RNF213 (right panel) at 4 hours post infection in HeLa cells. Data from *n=2* experiments, with error bars denoting the standard deviation.

### Both IpaH1.4 and IpaH2.5 bind and ubiquitylate RNF213

To identify the trans-acting factor antagonizing RNF213, we tested whether any of the secreted IpaH-family effectors from *S. flexneri* can bind to RNF213 in a pull-down assay using purified recombinant proteins. Out of all twelve effectors, only the virulence plasmid-encoded IpaH1.4 and IpaH2.5 proteins bound human RNF213 (**Figure 2A**). Cross-reactivity of IpaH1.4 and IpaH2.5 towards RNF213 is not unexpected as they are closely related proteins with identical leucine-rich repeats, the canonical substrate binding site in IpaH proteins **(Extended Data Figure 1A)**. To study the effect of IpaH1.4/2.5 on RNF213, we generated HeLa cells expressing GFP-tagged IpaH proteins or their catalytically inactive variants, in which the catalytically active cysteine in the NEL domain is replaced with an alanine to abolish E3 ubiquitin ligase activity. Expression of wild type IpaH1.4 or IpaH2.5, but not their catalytically inactive variants (IpaH1.4/2.5 C368A) or IpaH7.8, depleted RNF213 in the host cells (**Figure 2B**). Concomitantly, all active IpaH ligases but not their catalytically impaired variants induced their own degradation when over-expressed in mammalian cells (**Figure 2B**). To test the effect of IpaH1.4/2.5 on LPS ubiquitylation, we infected IpaH-expressing cells with *Salmonella* Typhimurium *ΔrfaL*, which is susceptible to LPS ubiquitylation and whose truncated LPS produces a characteristic banding pattern in the immunoblot for conjugated ubiquitin in heat-cleared bacterial lysate from infected cells^10^ (**Figure 2B**). Expression of IpaH7.8, which neither binds to nor depletes RNF213, had no effect on the ubiquitylation of LPS, while, in contrast, wild type IpaH1.4 or IpaH2.5 prevented LPS ubiquitylation, consistent with their ability to deplete RNF213. Catalytically inactive IpaH1.4/2.5 C368A did not prevent LPS ubiquitylation and did not reduce RNF213 protein levels, suggesting that mere binding of IpaH1.4 or IpaH2.5 to RNF213 is insufficient to inhibit RNF213 activity. We conclude that IpaH1.4 and IpaH2.5 antagonize LPS ubiquitylation through depletion of cellular RNF213 protein levels in a manner that requires IpaH E3 ubiquitin ligase activity.

**Figure 2.**
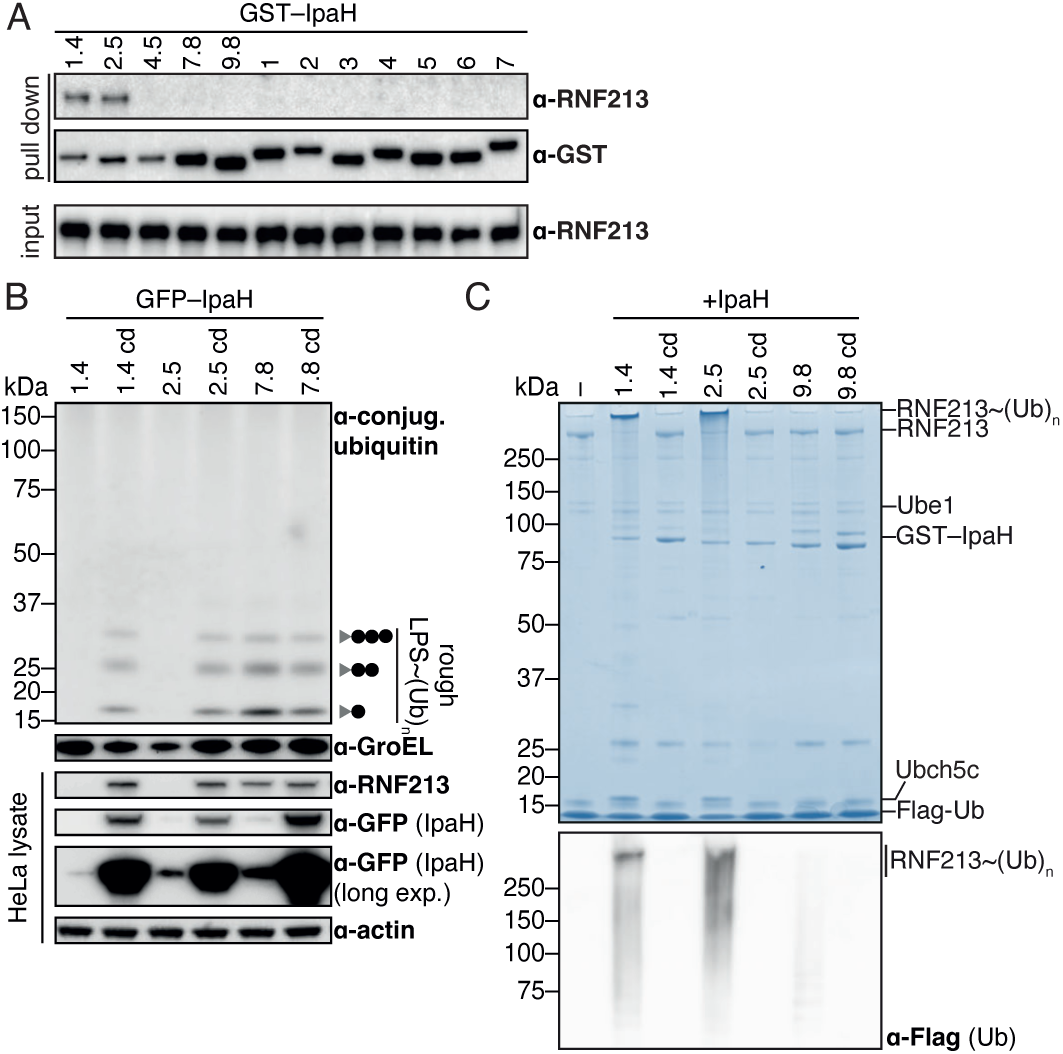
IpaH1.4 and IpaH2.5 bind to, ubiquitylate and antagonize RNF213. (A) HeLa cell lysate incubated with beads displaying the indicated GST-tagged IpaH proteins. Bound RNF213 detected by Western blot. (B) Immunoblot analysis of *S.* Typhimurium *ΔrfaL*, extracted from HeLa cells expressing the indicated GFP-tagged IpaH proteins. Grey triangle, *ΔrfaL* LPS; black circle, ubiquitin; GroEL, loading control for bacterial lysates; cd, catalytically dead (E3 inactive) variants; actin, loading control for HeLa lysates. The LPS fractions were isolated by heat clearance of bacterial lysates and probed for conjugated ubiquitin with FK2 antibody. The loading control (GroEL) was probed for in non-heat cleared bacterial lysates. All other proteins were probed for in host cell lysates. (C) Coomassie-stained gel of an *in vitro* ubiquitylation reaction containing Flag-tagged ubiquitin, UBE1, UBCH5C, the indicated IpaH proteins and enzymatically inactive RNF213 H4509A **(Extended Data Figure 2A**) as substrate. The majority of RNF213 is ubiquitylated by IpaH1.4 or 2.5, yielding a product of substantially increased molecular mass that migrates slower in the gel, as indicated. Lower panel, immunoblot for Flag-tagged ubiquitin.

To investigate the mechanism of RNF213 depletion by IpaH1.4 and IpaH2.5, we tested whether these proteins can ubiquitylate RNF213 *in vitro.* To avoid any unwanted background stemming from auto-ubiquitylation of RNF213, we used RNF213 H4509A, a catalytically inactive variant^10^, as substrate in the reaction **(Extended Data Figure 2A).** We found that in the presence of recombinant ubiquitin, E1 and E2 enzymes, RNF213 is ubiquitylated by either IpaH1.4 or IpaH2.5, but not by their catalytically inactive variants (IpaH1.4/2.5 C368A) or by IpaH9.8 (**Figure 2C, Extended Data Figure 2B)**. In summary, we discovered that IpaH1.4/2.5 can bind and ubiquitylate RNF213 *in vitro*, and, when recombinantly expressed in cells, target RNF213 for degradation, which prevents LPS ubiquitylation upon infection with *Salmonella* Typhimurium.

### IpaH1.4 protects *Shigella flexneri* from LPS ubiquitylation

We next tested whether the lack of RNF213-mediated LPS ubiquitylation in cells infected with *Shigella flexneri* is due to the action of IpaH1.4 and/or IpaH2.5 on RNF213. To this end we infected cells expressing GFP-RNF213 with Ruby-labeled *S. flexneri* wild type, *ΔipaH1.4* or *ΔipaH2.5* strains for analysis by flow cytometry (**Figure 3A, Extended Data Figure 3)**. When infected with either wild type bacteria or *S. flexneri ΔipaH2.5*, around 30% of the infected cells lost their GFP fluorescence at 4 hours post infection, indicative of RNF213 degradation. In contrast, upon infection with *S. flexneri ΔipaH1.4* no measurable RNF213 degradation occurred. Complementation of *S. flexneri ΔipaH1.4* with plasmid-encoded *IpaH1.4* under control of an inducible promoter fully recovered their ability to degrade RNF213 in infected cells (**Extended Data Figure 4A**). Endogenous RNF213 became similarly depleted in HeLa cells infected with *S. flexneri* wild type or *ΔipaH2.5* but not *ΔipaH1.4,* as assessed by either immuno-staining of endogenous RNF213 in infected cells or Western blotting of lysates from FACS-sorted infected cells **(Extended Data Figure 4B-C)**. We conclude that IpaH1.4, but not IpaH2.5, is required for RNF213 degradation by *S. flexneri.* To obtain further insight into the differential contribution of IpaH1.4 and IpaH2.5 to RNF213 degradation, we analyzed RNA extracted from cells infected with wild type *S. flexneri*, and confirmed that transcripts for both *IpaH1.4* and *IpaH2.5* were present, indicating that both genes are expressed and that likely a downstream mechanism, for example lack of IpaH2.5 protein secretion, limits the ability of IpaH2.5 to target RNF213 **(Extended Data Figure 5A)**. To investigate how ubiquitylated RNF213 is degraded, we infected GFP-RNF213 reporter cells with *S. flexneri* and treated them with either Carfilzomib or Bafilomycin A1, inhibitors of the proteasome or autophagy pathway, respectively (**Figure 3A, Extended Data Figure 3C)**. Only treatment with Carfilzomib prevented degradation of RNF213 upon infection with *Shigella*, indicating that IpaH1.4-induced degradation of RNF213 is proteasome-mediated, as is the case for other IpaH – host effector pairs.^34,36,39,40^

**Figure 3.**
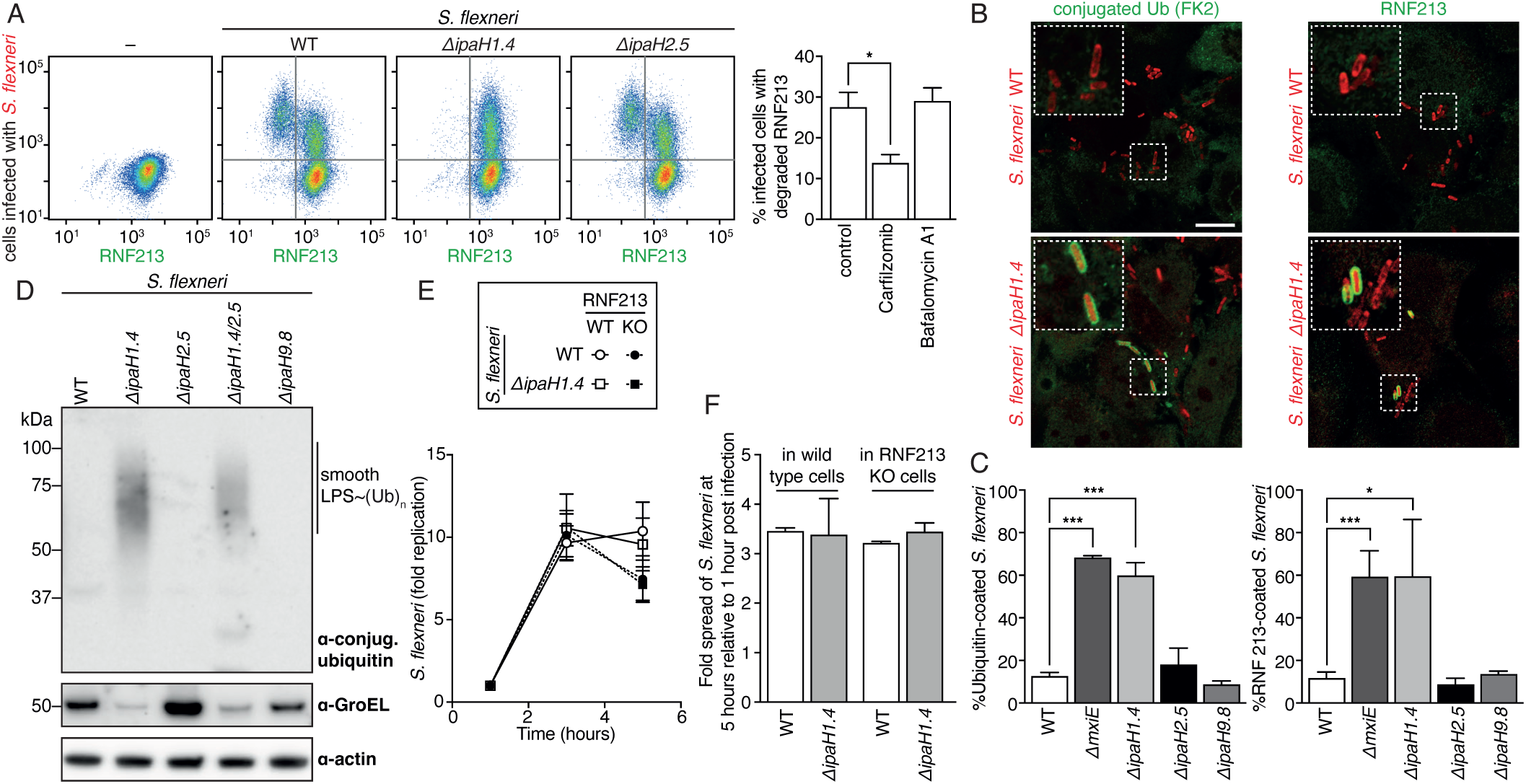
IpaH1.4-induced proteasomal degradation of RNF213 protects *Shigella* from LPS ubiquitylation. (A) Flow cytometry of GFP-RNF213 expressing MEFs, infected with the indicated strains of Ruby-labelled *S. flexneri* for 5 h. Quadrants depicted were used to quantify the percentage of infected cells in which degradation of RNF213 is observed. Note that cells with a higher bacterial burden are more likely to have lost RNF213. See also **Extended Data Figure 3A-B**. Bar graph: the fraction of infected cells, where RNF213 is degraded (upper left quadrant in **Extended Data Figure 3C**) in the presence of the indicated inhibitors. Data from *n*=4 experiments performed in duplicate. * p = 0.018, unpaired t-test. Error bars denote standard deviation. (B) Representative confocal micrographs of HeLa cells at 4 hours post infection with the indicated strains of Ruby-labeled *Shigella flexneri*, immunostained for conjugated ubiquitin (FK2) (left), or RNF213 (right). Dashed squares are magnified 2.5× in the insets. Scale bar, 10 μm. The corresponding micrographs of individual color channels are shown in **Extended Data Figure 5B**. (C) Percentage of the indicated *S. flexneri* strains positive for conjugated ubiquitin (detected by FK2 antibody) (left panel) and GFP-RNF213 (right panel) at 4 hours post infection in MEFs expressing human GFP-RNF213. Data from *n=*3 experiments, with error bars denoting the standard deviation. (D) Immunoblot analysis of the indicated strains of *S. flexneri*, which were extracted from MEFs at 4h post infection. The LPS fractions were isolated by heat clearance of bacterial lysates and probed for conjugated ubiquitin with FK2 antibody. The loading control (GroEL) was probed for in the non-heat cleared bacterial lysates. The host cell loading control (actin) was probed for in clarified host cell lysates. See also **Extended Data Figure 5C**. (E) Fold replication of the indicated strains of *S. flexneri* in wild type and RNF213 knock-out HeLa cells. Bacteria were counted based on their ability to grow on TSB plates. Proliferation is quantified from a single experiment, representative of two, including *n=6* technical replicates. Data are expressed as mean ± standard deviation. (F) Fold cell-to-cell spread of the indicated strains of *S. flexneri* in wild type and RNF213 knock-out HeLa cells at 5 hours post infection relative to 1 hour post infection, quantified from *n=3* experiments. Data are expressed as mean ± standard error in the mean.

We next tested whether the modulation of RNF213 protein levels by IpaH1.4 has functional consequences. We observed that knockout of IpaH1.4 allowed for recruitment of RNF213 to *Shigella* and rendered bacteria susceptible to RNF213-mediated ubiquitylation (**Figure 3B–D, Extended Data Figure 5B-C)**. In contrast, *S. flexneri ΔipaH2.5* was indistinguishable from the wild type strain in this experiment, confirming that IpaH1.4 is the sole IpaH-family effector antagonizing RNF213 under these conditions. Knockout of *mxiE*, a transcriptional regulator that stimulates expression of secreted effectors^42^, including the IpaH family, also rendered *Shigella* susceptible to coating with ubiquitin and RNF213 (**Figure 3C**). Next, we specifically tested the effect of IpaH1.4 on LPS ubiquitylation (**Figure 3D, Extended Data Figure 5C)**. We found that deletion of IpaH1.4, but not IpaH2.5 or IpaH9.8, rendered *Shigella flexneri* LPS susceptible to ubiquitylation. We conclude that IpaH1.4 causes proteasome-mediated degradation of RNF213 and thereby prevents LPS ubiquitylation, while IpaH2.5 does not contribute to the phenotype, at least in the context of *S. flexneri* infection in cell culture. Our observation is consistent with previous findings regarding IpaH1.4/2.5 and LUBAC, where, similarly, both effectors target LUBAC *in vitro*, but only IpaH1.4 affects LUBAC in infected cells.^40^

We then investigated whether RNF213-mediated LPS ubiquitylation restricts the proliferation or the cell-to-cell spread of *S. flexneri* lacking IpaH1.4. We found that *S. flexneri ΔipaH1.4* proliferated (**Figure 3E**) and spread (**Figure 3F**) in a manner comparable to wild type bacteria. This result indicates that RNF213-mediated ubiquitylation of bacterial LPS is insufficient to restrict *Shigella* proliferation or cell-to-cell spread, a perhaps surprising finding as RNF213 is crucial for restricting the proliferation of *Salmonella* Typhimurium in the host cytosol.^10^ It is therefore likely that *Shigella* has evolved additional strategies to block the downstream effects of RNF213-mediated ubiquitylation. These might include factors that block the recruitment of autophagy receptors and inhibit NF-κB activation, for example, IcsB, IpaH9.8, OspI and OspZ ^23,39,40,43–45^, or completely orthogonal means, such as actin-mediated motility, which may allow escape from the consequences of LPS ubiquitylation.

### IpaH1.4 binds the RING domain of RNF213

Finally, to gain molecular insights into the interaction between IpaH1.4/2.5 and RNF213, we determined the structures of human RNF213 bound to either IpaH1.4 or IpaH2.5 using cryogenic electron microscopy (cryoEM) (**Figure 4A-F, Extended Data Figures 6–7)**. The cryoEM maps show that the LRR domains of IpaH1.4 and IpaH2.5 interact with the RING domain of RNF213 (**Figure 4B–D, Extended Data Figure 6)**, whereas the unstructured N-termini and the catalytic NEL domains of IpaH1.4 and IpaH2.5 remain unresolved. Focused refinement of the RNF213– IpaH1.4/2.5 structure around the RING domain of RNF213 reveals molecular details of their interaction to 3.3 Å resolution (**Figure 4D**), a higher resolution reconstruction of the RNF213 RING domain than that previously achieved in published apo-structures of RNF213.^14,46^ IpaH1.4/2.5 interact exclusively with the RING domain of RNF213. Unsurprisingly, given the sequence identity of their LRR domains, IpaH1.4 and IpaH2.5 engage the RING domain of RNF213 in exactly the same way **(Extended Data Figure 6)**. Binding is mediated by the canonical concave substrate-binding side of the LRR, involving an extensive interaction interface spanning from strand β2 to β7. Several hydrophobic interactions (e.g., IpaH F120 / RNF213 F3992, IpaH V177/ RNF213 L4036) and hydrogen bonded amino acid pairs (e.g., IpaH K100 / RNF213 D4013, IpaH R215 / RNF213 Q4029, IpaH R157 / RNF213 C4035 carbonyl O) stabilize the interface (**Figure 4E, Extended Data Figure 8)**. Overall, the RNF213 RING domain docks into a net positively charged cavity in the LRR (**Figure 4F**) and into two hydrophobic patches formed by strands β6 and β8–10. Mutation of the key interfacial residues, including RNF213 L4036E and IpaH1.4 K100A, F120E, R215A, V217D, F238D abolish binding between RNF213 and IpaH1.4 **(Extended Data Figure 8)**. The flexible RNF213 loop between Gln4099 and Glu4112, which is not resolved in the RNF213 apo-structures^14,46^, nor in the IpaH1.4/2.5-bound structures reported here, must point towards the convex LRR face without forming a stable interaction. Focused refinements of the other domains of RNF213 revealed that the binding of IpaH1.4/2.5 does not cause any conformational changes in comparison to the previously published apo-structure of human RNF213.^14^ Although we did not resolve the NEL domain of IpaH1.4/2.5, locating the LRR at the RNF213 RING domain allows us to predict the region of RNF213 accessible to the ubiquitylating activity of IpaH1.4/2.5 (**Figure 4B**). Accounting for the length of the flexible linker between LRR and NEL domains in IpaH1.4/2.5, we predict that the NEL catalytic domain will have access to the CBM20 carbohydrate binding module of RNF213, its E3 module, in particular the M-lobe of the E3 shell, and possibly also to 32 lysine residues in the unstructured N-terminus of RNF213 (residues 1– 398), which is not resolved in any of the current cryoEM maps.

**Figure 4.**
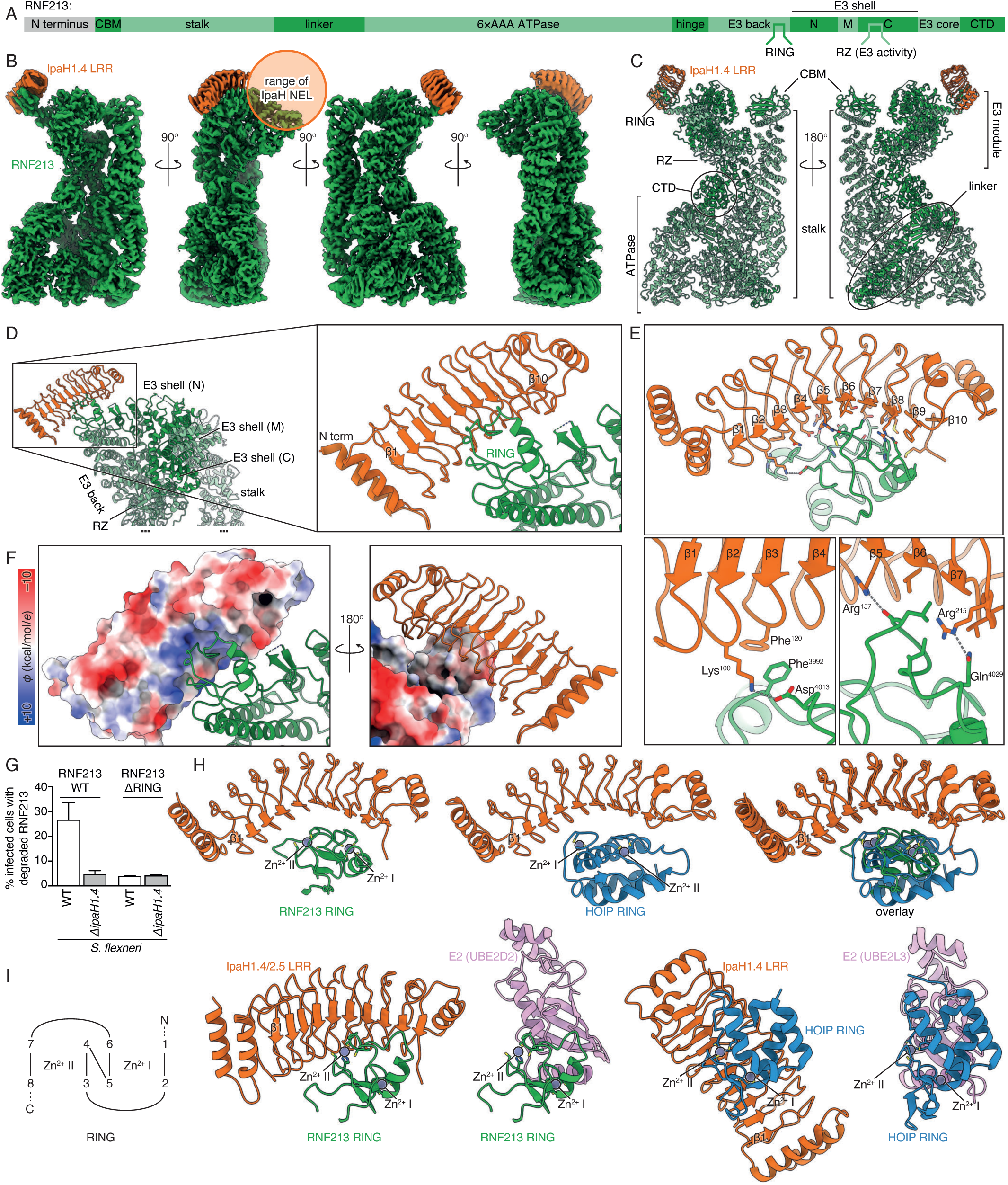
Structure of the IpaH1.4–RNF213 complex. (A) Domain annotation of human RNF213. (B) CryoEM map of the IpaH1.4–RNF213 complex at 2.9 – 3.3 Å resolution, and (C) corresponding atomic model, all colored by subunit: IpaH1.4, orange; RNF213, green. (D) Magnified view of the interaction interface between the IpaH1.4 LRR and the RNF213 RING domain. (E) Key interactions between IpaH1.4 and RNF213. (F) The interacting surface of IpaH1.4 with RNF213, shown from two opposing views, colored by electrostatic potential, *φ*. (G) Degradation of RNF213 and RNF213 ΔRING by *S. flexneri* WT and *ΔipaH1.4*, quantified by flow cytometry **(Extended Data Figure 3D**). (H) Structure comparison between IpaH1.4 LRR–RNF213 RING (this work) and IpaH1.4 LRR–HOIP RING1 (PDB 7V8G), aligned by the IpaH1.4 LRR. (I) Structure comparison between IpaH1.4 LRR–RNF213 RING (this work), UBE2D2–RNF213 RING (predicted), IpaH1.4 LRR–HOIP RING1 and UBE2D2–HOIP RING1 (PDB 7V8G & 7V8F)^41^, all aligned by the Zn fingers in the RING domains, and colored by subunit: IpaH1.4, orange; RNF213, green; UBE2D2, pink; HOIP, blue. The Zn fingers are labeled I and II as per the shown domain architecture diagram (1–8 denote the Zn coordinating residues, Cys or His).

The cryoEM model predicts that deletion of the RNF213 RING domain abolishes the interaction between RNF213 and IpaH1.4/2.5. Indeed, in a flow-cytometry based cellular assay (**Figure 4G, Extended Data Figure 3D**) we observe that RNF213 ΔRING is no longer degraded in cells infected with *S. flexneri*, which confirms that the RING domain is the relevant binding site in cells. Since the catalytic activity of RNF213 towards LPS is mediated by the RZ domain^10^, binding of IpaH1.4/2.5 to the distal RING domain explains why binding alone does not inhibit the activity of RNF213 (**Figure 2B**).

Interestingly, IpaH1.4 targets at least two enzymes (RNF213 and the LUBAC subunit HOIP), which are involved sequentially in the anti-bacterial ubiquitylation cascade.^10,11,40^ Since IpaH1.4 recognizes both HOIP and RNF213 via their RING domains (**Figure 4H** and ^41^), the question arises whether IpaH1.4 may target additional RING domain containing proteins and how specificity is conferred. We compared the structures of IpaH1.4 bound to the RING1 domain of the LUBAC subunit HOIP (PDB 7V8G)^41^ and the RING domain of RNF213 reported here by superimposing IpaH1.4 (**Figure 4H**) or the RING domains (**Figure 4I**). This comparison reveals that binding of IpaH1.4 to RNF213 and HOIP is mutually exclusive and that in either complex IpaH1.4 engages and occludes the E2-binding face of the RING domains. However, the respective RING domains bind the LRR in different orientations, with the Zn^2+^ site I and site II of the RNF213 RING domain aligning with sites II and I of the LUBAC RING1 domain, respectively, when the structures of the two complexes are aligned by the IpaH1.4 LRR (**Figure 4H**). These two binding modes are mediated through interactions with distinct LRR residues so that only IpaH1.4 Lys100 and Arg157 appear engaged with both RING substrates (**Extended Data Figure 8B**),^41^ while the residues with which IpahH1.4 Lys100 and Arg157 interact are not topologically equivalent between the two RING domains. We conclude that the distinct binding mode of the HOIP and RNF213 RING domains reveals significant substrate specificity in IpaH1.4/2.5, a finding that argues against the possibility of IpaH1.4/2.5 globally targeting RING domain proteins.

However, we nevertheless expect that IpaH1.4 and IpaH2.5 might be able to engage select additional RING E3 ligases via their E2-binding interfaces. We therefore performed an unbiased pull-down mass spectrometry screen for IpaH1.4 and IpaH2.5 interactors from HeLa cell lysates (**Figure 5A, Supplementary Table 3).** We also tested lysates from interferon-γ treated cells, since IpaH-family effectors are known to associate with certain interferon-induced host proteins, such as guanylate-binding proteins (GBPs).^33,34^ In addition to RNF213 and HOIP, we found six other RING-domain containing proteins specifically enriched in the IpaH1.4 and/or IpaH2.5 pulldowns, namely TRIM25, TRIM29, TRIM47, DTX3L, TRAF2, BIRC3, which, remarkably, have all been implicated in inflammatory signaling and immunity. Whether and how any of these proteins are targeted by the *Shigella* effectors during infection remains to be determined. Computational structure predictions position the IpaH1.4/2.4 LRR domain at the E2-interacting interface of the RING domains of each potential interactor, although the relative orientation of the domains varies between individual complexes (**Figure 5B–C**). We conclude that the *Shigella* effectors IpaH1.4/2.5 may have evolved to target a limited number of select RING E3 ligases involved in immune signaling, in addition to RNF213 and LUBAC, as part of the *Shigella* strategy to manipulate host immunity.

**Figure 5.**
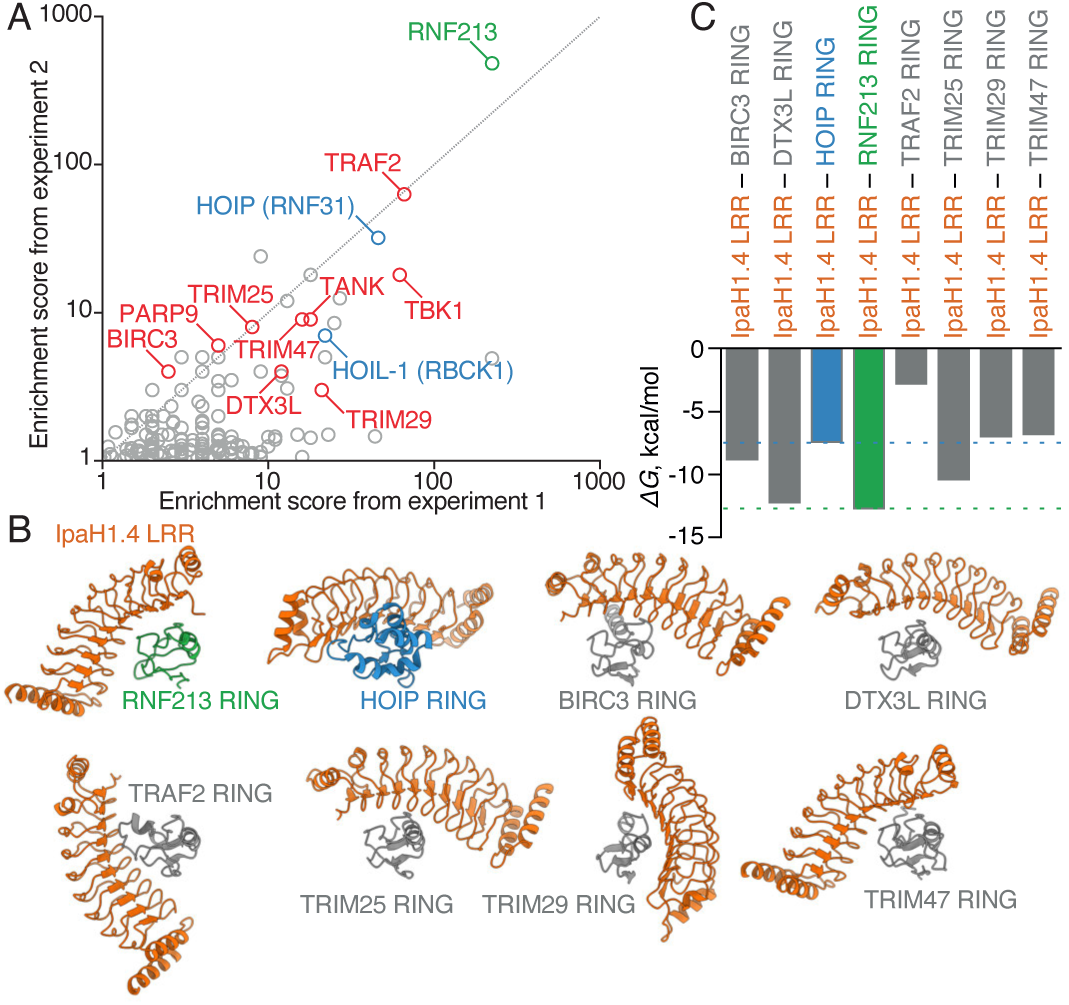
Mass spectrometry-based identification of potential RING-domain containing targets of IpaH1.4/2.5 in human cells. (A) Truncated scatter plot of proteins enriched in pull downs with GST-IpaH1.4 and IpaH2.5 relative to negative controls (GST and GST-IpaH9.8). The enrichment score is defined as the ratio (summed number of peptides detected in the IpaH1.4 and IpaH2.5 samples + 1) / (summed number of peptides detected in the control samples + 1). Enrichment scores from two separate experiments are plotted; only proteins with an enrichment score >1.0 in both experiments are shown. For a complete list see **Supplementary Table 3**. RNF213 is highlighted in green, LUBAC subunits HOIL-1L and HOIP in blue, other RING-domain containing proteins and their known binding partners in red, e.g. PARP9/DTX3L^72^ and TRAF2/TBK1/TANK.^73^ (B) Models of the IpaH1.4 LRR interaction with RING domains of RNF213 and HOIP (experimental); BIRC3, DTX3L, TRAF2, TRIM25, TRIM29 and TRIM47 (predicted). The predicted models were obtained from full-length protein sequences using AlphaFold 3. Only LRR and RING domains are shown for clarity. All structures are displayed with their RING domains in identical orientation. (C) Binding energies, *ΔG*, for experimentally determined (colored) and predicted (grey) interfaces, calculated using PDB PISA.^74^ Dashed lines, *ΔG* range for the two experimentally determined interfaces (IpaH1.4 LRR with RNF213 RING and with HOIP RING). Three of the six predicted interfaces have binding energies within this range.

## Discussion

Here we report that *S. flexneri* uses the secreted effector IpaH1.4 to bind, ubiquitylate and target the host E3 ligase RNF213 for degradation via the proteasome, thereby evading LPS ubiquitylation in the cytosol of infected host cells. In the ensuing arms race between pathogen and host, the host seeks to ubiquitylate and target the pathogen for degradation, while the pathogen aims to ubiquitylate and degrade the host ubiquitylation machinery. IpaH1.4 represents the first known example of a protein from a cytosol-dwelling pathogen to directly antagonize RNF213-mediated LPS ubiquitylation. In contrast to such direct antagonism by IpaH1.4, *Burkholderia thailandensis* and certain *Chromobacterium* species deploy the deubiquitylase TssM, to reverse, rather than prevent, ubiquitylation of LPS.^47^ Yet another bacterial anti-RNF213 strategy is exemplified by the *Chlamydia trachomatis* effector GarD, a putative transmembrane protein, which prevents RNF213 recruitment to the pathogen-containing vacuole in *cis*, but its mechanism remains to be understood.^16^ The fact that RNF213 is targeted directly and indirectly by bacterial effector proteins from distinct species and through a variety of strategies highlights the evolutionary importance of RNF213 in restricting intracellular pathogens. Whether further strategies against RNF213-mediated ubiquitylation have evolved in other cytosol-dwelling pathogens should be investigated as it may aid in elucidating the molecular mechanisms of RNF213 and in the development of novel therapeutic strategies.

Coincidentally, both RNF213 and IpaH1.4 are unconventional E3 ubiquitin ligases with poorly understood catalytic mechanisms. The E3 ligase activity of RNF213 and IpaH-family effectors are found in their RZ and NEL domains, respectively.^10,26^ The structural basis for trans-thiolation catalysis by either of these domains has not been visualized yet. In the structure of IpaH1.4/2.5 in complex with RNF213 presented here, we did not resolve the NEL domain of the bacterial ligase, despite extensive classification and local refinements, which is likely due to the flexible linker between the LRR and the NEL, and is entirely consistent with the propensity of the NEL domain to engage with a multitude of nearby ubiquitin acceptors.

Interestingly, IpaH1.4 antagonizes anti-bacterial ubiquitylation on multiple levels. In addition to preventing LPS ubiquitylation through RNF213 degradation as reported here, IpaH1.4 also blocks downstream M1-linked ubiquitin chain formation by LUBAC in an analogous manner, i.e. through binding the E2-interacting face of the HOIP RING1 domain and subsequent ubiquitylation of the host ligase.^11,40^ We have previously shown that, at least in the context of cytosolic *Salmonella*, LPS ubiquitylation by RNF213 is required for the recruitment of LUBAC to the pathogen and further M1-linked ubiquitin chain formation.^11^ Therefore, a single bacterial effector, IpaH1.4, has evolved to target multiple host enzymes that are all part of the same anti-bacterial ubiquitylation pathway. It even seems that not only RNF213 and LUBAC but also further immune-related RING E3 ligases are targeted by the *Shigella* effector IpaH1.4. The putative candidates, shown in **Figure 5A**, include the ADP-ribosylation complex DTX3L/PARP9 with the tantalizing ability to ubiquitylate nucleic acids and ADP-ribosylated substrates *in vitro*^48–50^, and the ISG15 E3-ligase TRIM25^51^, among others. Their interactions with the *Shigella-*secreted effectors remain to be characterized further and may offer new insights into their roles in anti-bacterial innate immunity. Nevertheless, deletion of IpaH1.4, either alone or in combination with IpaH2.5, was insufficient for anti-bacterial ubiquitylation to restrict *Shigella* replication, a situation reminiscent of TssM in *Burkholderia thailandensis*, whose deletion also did not affect bacterial replication in response to anti-bacterial ubiquitylation.^47^ Such lack of phenotype is often due to the presence of additional inhibitors, particularly in pathogens with a plethora of effectors, such as *Legionella pneumophila*, but is also known from *Shigella*, where host pathways are simultaneously targeted by multiple effectors, for example IpaH9.8 and OspC3, which inhibit GBPs and caspase-4, respectively.^34,52^

RNF213 is unusual in harboring two putative E3 ligase domains: an RZ finger, whose E3 ubiquitin ligase function is well established, and a RING domain, whose function remains elusive. The RNF213 RING domain is atypical in that it lacks the ‘linchpin’ Arg/Lys residue normally required to catalyze ubiquitin transfer from a charged E2 enzyme^53^ and it appears to contribute neither to the auto-ubiquitylating activity of RNF213 *in vitro* **(Extended Data Figure 2A)** nor to the ubiquitylation of LPS by RNF213.^10^ However, missense mutations in the RNF213 RING finger in patients with Moyamoya disease strongly suggest functional importance to the domain.^54,55^ We anticipate that the IpaH1.4 LRR domain may become a useful biochemical tool to further the study of the RNF213 RING domain. The fact that the IpaH1.4-binding interface of the RING domain, despite the evolutionary pressure exerted by *Shigella* IpaH1.4, has not evolved away, suggests that the RNF213 RING has some important function and that binding partners in the host cell may remain to be discovered.

## Methods

### Plasmids, antibodies & reagents

The open reading frames encoding IpaH-family proteins were cloned into M6P plasmids as N-terminal GFP fusions; these were used to produce recombinant murine leukaemia viruses for stable protein expression in mammalian cells.^56^ For recombinant protein expression in E. coli, the same genes were cloned into pETM-30 plasmids. Catalytically inactive (cd) variants of the IpaH proteins were produced by mutating the catalytic cysteine to alanine, as previously described.^34^ Briefly, the following mutant IpaH proteins were encoded as catalytically inactive variants: *ipaH1.4* C368A, *ipaH2.5* C368A, *ipaH7.8* C357A, *ipaH9.8* C337A.

The following primary antibodies were used in this work: FK2 (Enzo Life Science, BML-PW8810, against conjugated ubiquitin), anti-GroEL (Enzo Life Science, ADI-SPS-875-F), anti-actin (Abcam, ab8227), anti-RNF213 (Merck, HPA003347 and HPA026790), anti-Flag–M2–HRP (Merck, A8592), anti-GFP (JL8, Clontech, 632381), anti-K48-linked ubiquitin (Abcam, ab140601), anti-*Salmonella* typhimurium LPS (BioRad, 8210-0407), *Salmonella* O Antisera Group B (BD Difco, 229481). The secondary antibodies used were from Thermo Fisher Scientific (Alexa-conjugated anti-mouse, anti-goat, anti-human, and anti-rabbit antisera) and Dabco (HRP-conjugated reagents).

### Cell culture

HeLa, 293ET and mouse embryonic fibroblasts (MEFs) were grown in IMDM supplemented with 10% FCS at 37°C in 5% CO2. Cell lines have not been authenticated. All cell lines were tested and found to be negative for *Mycoplasma*. Stable cell lines were generated by retroviral transduction, except for the Flag-GFP-RNF213 lines, which were generated using an inducible PiggyBac transposon system, as previously described.^10^ The *RNF213*-knockout cell lines used were also previously described.^10^ HeLa cells, recombinantly over-expressing IpaH proteins (wild-type and catalytic-dead variants) were generated by viral transduction, followed by antibiotic selection.

### Bacteria

*Escherichia coli* strains MC1601 and NEB 10-beta were used for plasmid production and the BL21(DE3) strain was used for protein purification. All *E. coli* strains were grown on tryptic yeast extract (TYE) agar plates or in Luria broth (LB) or Terrific broth (TB). *Salmonella enterica* serovar Typhimurium strain 12023 and the isogenic *ΔrfaL* were provided by D. Holden. *S.* Typhimurium was grown in LB or on LB agar plates. *Shigella flexneri* strain M90T (wt and *ΔrfaL)* was provided by Chris Tang (University of Oxford), whereas strains M90T *ΔmxiE*^57^ and *ΔipaH9.8*^57^ and *ΔipaH1.4*, *ΔipaH2.5* and *ΔipaH1.4/2.5*^40^ together with isogenic wild type controls were provided by John Rohde (Dalhousie University, Nova Scotia, Canada) and Neal Alto (University of Texas Southwestern Medical Center, US), respectively. *S. flexneri* was grown in tryptic soy broth (TSB) or on tryptic soy agar containing 0.003% Congo red.

### Bacterial infections

The indicated strains of *S.* Typhimurium were grown overnight in LB and subcultured (1:33) in fresh LB for 3.5 hours before infection. The indicated strains of *S. flexneri* were grown overnight in TSB and subcultured (1:100) in fresh TSB for 2.5 hours before infection. Host cells for infection were seeded in 24-well or 6-well plates, or 10 cm dishes, and cultured in antibiotic-free media (IMDM/10% FCS) for one hour prior to infection. To infect cells with *S.* Typhimurium, 500 μL of bacterial subculture were added per 10 cm dish, or the linearly scaled amount for cells in multi-well plates. To infect cells with *S. flexneri*, the bacterial subculture was first washed in PBS, then re-suspended in antibiotic-free cell media without dilution, 2 mL of the bacterial suspension were added per 10 cm dish, and the dishes were centrifuged at 845g for 10 minutes at room temperature. The infected cells, in both cases, were incubated at 37°C for 30 minutes. After that, the cells were washed two times with warm PBS and cultured in 100 μg/mL gentamycin (after *S.* Typhimurium infection) or 50 μg/mL gentamycin (after *S. flexneri* infection). Infected cells were harvested 4 hours post infection. The cells were washed once with PBS and lysed in 1 mL lysis buffer (PBS, 0.1% Triton X-100, 2 mM iodoacetamide, protease inhibitors) at 4°C. The cell lysates were centrifuged at 400g for 5 min at 4°C and the supernatant containing the post-nuclear cell lysate and the bacteria was collected and centrifuged at 16,100g for 10 min at 4°C. The supernatant, containing the clarified host cell lysate, was saved, and the bacterial pellet was washed once with PBS, followed by bacterial lysis in 35 μL BugBuster (Merck) including 2 mM iodoacetamide for 5 minutes at room temperature with occasional vortexing. The bacterial lysate was centrifuged at 16,100g for 10 min at room temperature, and the supernatant (clarified bacterial lysate) was split up in 2 fractions. One part was directly mixed with Laemmli buffer (Bio-Rad) containing 100 mM DTT and boiled for 5 min. The remainder of the clarified bacterial lysate was used to further purify ubiquitylated LPS. It was heated at 100°C for 15 min, centrifuged at 16,100g for 10 min at room temperature, and this heat-cleared supernatant was mixed with Laemmli buffer containing 100 mM DTT and boiled for 5 min.

### Immunoblot analysis

Post-nuclear supernatants from HeLa or MEF cells, bacterial lysates from these infected cells, and bacterial LPS from these infected cells were obtained as described above. Heat-cleared bacterial lysates were used to visualize ubiquitylated LPS with FK2 antibody; non-heat-cleared bacterial lysates were used to probe for GroEL as a loading control; post-nuclear HeLa or MEF lysates were used to probe for host proteins. All samples were mixed with Laemmli buffer containing 100 mM DTT and boiled for 5 min. Samples were run on NuPAGE 4–12% Bis-Tris gels (Thermo Fisher Scientific) in MOPS SDS running buffer at 180 V for 70 min. Overnight wet transfer at 20 V was used to transfer proteins onto methanol-activated PVDF membranes (Millipore). Membranes were blocked for 1 hour in PBS-T (137 mM NaCl, 2.7 mM KCl, 10 mM Na2HPO4, 1.8 mM KH2PO4, 0.1% Tween 20) containing 5% bovine serum albumin (BSA), followed by overnight incubation at 4°C or 1 – 4 hour incubation at room temperature with primary antibodies in PBS-T containing 5% BSA. Membranes were washed in PBS-T, incubated with secondary HRP-conjugated antibodies in PBS-T containing 5% milk for 1 hour at room temperature, followed by three PBS-T washes. Visualization following immunoblotting was performed using ECL detection reagents (Amersham Bioscience) and a ChemiDoc MP imaging system (Bio-Rad). Alternatively, instead of HRP-conjugated antibodies, fluorophore-conjugated antibodies were used in some instances, in which case the blots were visualized directly after the final wash.

### Microscopy of fixed cells

Cells were grown on glass coverslips, and after infection and fixation in 4% paraformaldehyde were washed twice in PBS, permeabilized for 5 min in PBS with 0.1% Triton X-100, and blocked in PBS with 2% BSA for 1 hour. Cells were incubated overnight with primary antibodies followed by Alexa-conjugated secondary antibodies in blocking solution for at least 1 hour at room temperature. Coverslips were then mounted in Prolong gold mounting medium. Images of infected cells were acquired on a Zeiss 780 inverted microscope using a 63x/1.4NA oil immersion lens. Images were analyzed using Fiji.^58^ Marker-positive bacteria were scored by eye from coverslips on a Zeiss Axio Imager microscope using a 100x/1.4NA oil immersion lens, amongst at least 200 bacteria per experiment. In the case of *S*. Typhimurium, only large (cytosolic) bacteria were considered for quantification.

### Enumeration of intracellular bacteria (CFU assays)

A slight variation on the above protocol for bacterial infection was used to infect cells for enumeration of intracellular bacteria. *S. flexneri* was grown overnight in TSB and sub-cultured (1:100) in fresh TSB for 2.5 hours before infection. Such cultures were consecutively washed in PBS and re-suspended in antibiotic-free IMDM/10% FCS immediately before 10 μL was used to infect HeLa or 20 μL for MEF cells in 24-well plates. The plates with infected cells were centrifuged for 10 min at 670g followed by incubation at 37°C for 15 min. Following two washes with warm PBS, cells were cultured in 100 μg/mL gentamycin for 1 hour and 20 μg/mL gentamycin thereafter. To enumerate intracellular bacteria, cells from triplicate wells were lysed in 1 mL PBS containing 0.1% Triton X-100. Serial dilutions were performed in PBS and plated in duplicates on TYE agar.

### Flow cytometry

A slight variation on the above protocol for bacterial infection was used to infect cells for analysis by flow cytometry. MEF cells harboring expression cassettes for doxycycline-inducible GFP-RNF213^10^ or GFP alone were cultured on 24-well plates for 48 hours prior to infection with the indicated strains of *S. flexneri*. GFP-RNF213 or GFP expression was induced by the addition of 1 μM doxycycline to antibiotic-free media (IMDM/10% FCS) 18 hours prior to infection. The cells were washed twice in warm PBS and infected with 100 μL *S. flexneri* in IMDM/10% FCS for 30 min in triplicate wells, as described above. To test the effects of autophagy and proteasome inhibitors, the cells were treated with 100 nM Bafilomycin A1 or Carfilzomib, respectively, or DMSO as a negative control. The ligands were added 15 minutes pre-infection and maintained throughout the experiment. Cells were cultured in 100 μg/mL gentamycin for 1 hour after infection and 20 μg/mL gentamycin thereafter. When testing the GFP-RNF213 ΔRING variant, the cells were also treated with the pan-caspase inhibitor Q-VD (20 μM) at the time of doxycycline induction to prevent RNF213 ΔRING-induced cell death. Q-VD was removed prior to infection. For the complementation of *S. flexneri ΔipaH1.4*, full length *ipaH1.4* was cloned into the plasmid p4928 (Addgene #168177, gift from Michael Hensel).^59^ Expression of ipaH1.4 was induced by addition of 100 ng/mL Anhydrotetracycline (ATc) to the *S. flexneri* subculture 1 h before infection and maintained throughout the experiment. Cells were washed with PBS and trypsinized at either 1 h or 5 h post infection, centrifuged at 400g for 2 min and washed twice with PBS. Cells were fixed in 4% paraformaldehyde for 15 min, washed twice in PBS and resuspended in PBS/1% BSA.

For detection of endogenous RNF213 by flow cytometry Hela WT or RNF213 KO cells were infected with the indicated strains of *Shigella flexneri* for 30 min and trypsinized at 4 h post-infection. Cells were pelleted at 400g, washed once in PBS and subsequently fixed and permeabilized using Cell Fixation and Permeabilization Kit (Nordic MUbio) – anti-RNF213 antibody (HPA026790) was included during permeabilization for 45 min. Cells were washed twice in PBS/5% fetal calf serum and incubated with Alexa-fluor 488-conjugated anti-rabbit secondary antibody at 1:500 dilution in permeabilization buffer for 30 min. Cells were washed twice and resuspended in PBS/5% FCS. All samples were run on a LSRFortessa flow cytometer, collecting a total of 30,000 events per sample. The quantitative difference in staining of RNF213 WT and KO cells was interpreted as the specific contribution of RNF213 to the assay.

For bacterial spreading experiments cells were infected with a relatively low MOI of Ruby-expressing *S. flexneri* in order to achieve 2–5% infected cells at 1 h post-infection as assessed by flow cytometry. This would presumably permit each individual infected cell to spread *S. flexneri* to neighboring uninfected cells. Infection was allowed to proceed for 5 h at which point the percentage of infected cells was quantified and the fold spread was calculated as (% infected at 5 h)/(% infected at 1 h). All data were analyzed in FlowJo (BD Biosciences).

### Flow-activated cell sorting

Live, rather than fixed, infected cells were sorted using a BD FACSAria Fusion flow-activated cell sorter. HeLa cells were infected with the indicated strains of *Shigella flexneri* as described in the section ‘Bacterial infections’ above. The cells were harvested by scraping into PBS at 4 hours post infection and washed twice with PBS. The cells were sorted into three populations, based on the *Shigella-*encoded mRuby fluorescence: uninfected, low fluorescence, and high fluorescence, where the fluorescence intensity of each cell serves as a proxy for bacterial load in the cell. For each specimen 10,000 cells were sorted into each of the three populations. The collected cells were pelleted by centrifugation and resuspended in 10 μL lysis buffer (8 M urea, 50 mM Tris-HCl, pH 8, 0.5% NP40, 0.1% SDS), supplemented with 1:100 universal nuclease (Pierce). The cell lysates were mixed with Laemmli buffer, boiled, and subjected to immunoblot analysis as described above.

### Detection of IpaH1.4/2.5 transcripts in infected cells

MEF RNF213 KO cells were infected with *S. flexneri* for 30 minutes, as described above, washed twice in warm PBS and cultured for a further 30 min. Cells were lysed in Buffer RLT (Qiagen) with the addition of 143 mM β-mercaptoethanol, and RNA was extracted using an RNeasy Extraction Kit (Qiagen). cDNA was synthesized using Superscript III Reverse Transcriptase (Thermo Fisher) and random hexamer primers from 1.5 μg RNA per sample. Real-time quantitative Taqman PCR was performed using primers specific for *ipaH1.4* or *ipaH2.5* and FAM-MGB-labelled probes as listed in **Supplementary Table S2**. Primers and probe for the murine *Actb* gene (Assay ID Mm01205647_g1, Thermo Fisher) were used to normalize data across samples. Plasmids harboring *ipaH1.4* or *ipaH2.5* were used to quantify the number of molecules of each per infected well amounting to approximately 10^5^ cells. Cells infected with either *S. flexneri ΔipaH1.4* or *ΔipaH2.5* were used to demonstrate the specificity of the primers and probes for either gene.

### Protein production in *E. coli*

IpaH1.4/2.5 were recombinantly over-expressed in *E. coli* BL21(DE3) as 6×His-GST N-terminal fusions, both in the pETM-30 vector. The plasmids were transformed into BL21(DE3) cells. Overnight cultures, started from a single colony, were used to inoculate 2 L of TB (supplemented with 50 μg/mL kanamycin) at a ratio of 1:200. The cultures were grown at 37°C and 200 rpm until reaching an OD at 600 nm of 2 and then induced with 1 mM IPTG for 20 h at 200 rpm, 18°C. Following collection by centrifugation, the cell pellets were resuspended in 200 mL lysis buffer (20 mM Tris-HCl pH 8, 300 mM NaCl), supplemented with 0.5 mM TCEP, protease inhibitor (cOmplete EDTA-free tablets, Roche), lysozyme (1 mg/mL, Sigma), and 5μL universal nuclease (Pierce). Cells were disrupted using a high-pressure homogenizer at 38,000 psi and 4°C. Cell debris were removed by centrifugation at 30,000g for 30 min at 4°C, and the volume of the supernatants was adjusted to 350 mL each by adding lysis buffer. Proteins were purified by immobilized metal-affinity chromatography. The cell lysates were mixed with 2.5 mL Ni-NTA Agarose resin (Qiagen). Binding to the resin was allowed to proceed for 2 hours while shaking at 110 rpm at 10°C. The beads were then collected by gravity flow and washed three times with 50 mL of wash buffer (25 mM Tris-HCl pH 8, 400 mM NaCl, and 0.5 mM TCEP), containing increasing imidazole concentrations: 25 mM, 37.5 mM, and 50 mM, respectively. The proteins were eluted in 20 mL of elution buffer (25 mM Tris-HCl pH 8, 400 mM NaCl, 0.5 mM TCEP, 500 mM imidazole), and concentrated to around 2 mL final volume. The 6×His-GST tag was removed by incubation with His-tagged TEV protease at 4°C for 13 hours in cleavage buffer (50 mM Tris-HCl pH 8, 150 mM NaCl, 0.5 mM EDTA, 0.5 mM TCEP). Untagged IpaH1.4/2.5 were further purified either by reverse immobilized metal-affinity chromatography (for use in cryoEM) or by anion exchange chromatography using a Resource Q column (Cytiva), in buffer 20 mM Tris-HCl pH 8.5, 10 – 500 mM NaCl, 4 mM DTT, and size exclusion chromatography using a Superdex 200 16/60 column (Cytiva), the latter in buffer 20 mM Tris-HCl pH 8.5, 200 mM NaCl, 4 mM DTT (for all other experiments).

### Protein production in *S. frugiperda*

Wild-type human RNF213 was expressed in insect cells and purified as previously described.^10,14^ Briefly, pOP806_pACEBac1 2xStrep-RNF213 plasmid was transformed into DH10EmBacY cells (DH10Bac with YFP reporter). Blue–white screening was used to isolate colonies containing recombinant baculoviral shuttle vectors (bacmids) and bacmid DNA was extracted combining cell lysis and neutralization using buffer P1, P2 and N3 (Qiagen), followed by isopropanol precipitation. A 6-well plate of *Spodoptera frugiperda* (Sf9) cells (Oxford Expression Technologies) grown at 27°C in Insect-Xpress media (Lonza) without shaking was transfected with bacmid plasmid using PEI transfection reagent. After 6 days, virus P1 was collected and used 1:25 to transduce 50 mL (1.8 × 10^6^ cells/ml) of Sf9 cells. After 7 days of incubation at 27°C with 140 rpm shaking, virus P2 was collected. To express protein, 1 L (2.6 × 10^6^ cells/mL) of Sf9 cells was transduced with 1:50 dilution of P2 virus and incubated at 27°C with 140 rpm shaking for 72 hours. Cells were pelleted by centrifugation, snap-frozen in liquid nitrogen, and stored at −80°C. To lyse cells, the pellets were thawed and resuspended to 180 mL total volume in lysis buffer (30 mM HEPES, 100 mM NaCl, 10 mM MgCl2, 0.5 mM TCEP, pH 7.6), containing 20 μL universal nuclease (Pierce), 20 μL benzonase nuclease (Sigma Aldrich), and 2 EDTA-free protease inhibitor tablets (Roche). The cell suspension was stirred for 1 hour and then sonicated for 50 seconds in 5-second pulses with 25-second waiting time at 70% amplitude using a 130 W microtip sonicator (Vibra Cell). The lysate was centrifuged at 20,000g for 60 min at 4°C. The clarified lysate was filtered through a 0.2 μm filter (Millipore) and applied to 2 × 5-mL StrepTrap HP columns (GE Healthcare), connected in series. After washing the columns with 100 mL lysis buffer, RNF213 was eluted with lysis buffer supplemented with 2.5 mM desthiobiotin, pH 8. The eluted protein was kept at 4°C and used immediately for cryoEM specimen preparation. All purification steps were carried out at 4°C, using an ÄKTA pure 25 (GE Healthcare). RNF213 variants, including RNF213 ΔRING, RNF213 H4509A and RNF213 E2488A/E2845A, were expressed and purified analogously.

### *In vitro* ubiquitylation assays

First, pre-charged E2 enzyme was prepared by incubating 10 μM UBCH5C (Boston Biochem) with 200 μM FLAG-ubiquitin (Boston Biochem) and 0.2 μM UBE1 in reaction buffer (30 mM HEPES pH7.4, 100 mM NaCl, 10 mM MgCl2, 50 mM ATP, 5 mM DTT) for 15 minutes at 37°C. UBE1 was purified as previously described.^10^ The pre-charged E2 reaction was diluted five-fold with the addition of 350 nM recombinantly produced RNF213 (for RNF213 auto-ubiquitylation experiments) or 350 nM recombinantly produced RNF213 H4509A and 2 μM recombinantly produced IpaH proteins (for IpaH-mediated ubiquitylation of RNF213). The reaction was incubated at 37°C for an additional hour, then further processed for immunoblot analysis or directly visualized in the gel by Coomassie staining.

### Pull down assays

Recombinantly expressed catalytically dead IpaH proteins were purified from *E. coli* using a slight variation on the above procedure. Briefly, *E. coli* carrying the GST-IpaH (catalytically dead cysteine to alanine mutant) constructs in a pETM30 plasmid were grown to optical density at 600 nm of 0.7, and protein expression was induced with 0.1 mM of IPTG for 20 hours at 200 rpm, 18°C. Following collection by centrifugation, the cell pellets were resuspended in 10 mL lysis buffer per 1 L of bacterial culture (20 mM Tris-HCl pH 8, 150 mM NaCl, 10 mM MgCl2), supplemented with 5 mM TCEP, protease inhibitor (cOmplete EDTA-free tablets, Roche), 20 mM imidazole, and 20 μg/mL DNase (Sigma). Cells were disrupted by freeze-thawing, followed by high-pressure homogenization at 38,000 psi and 4°C. Cell debris were removed by centrifugation at 30,000g for 30 min at 4°C, and the supernatants were bound to pre-equilibrated GST-Sepharose resin (Cytiva, 1 mL per 1 L bacterial culture) in presence of 0.1% Triton X-100 for 1 hour at 4 °C. The resin was washed in 10 mL lysis buffer five times, and the GST-tagged proteins were eluted in 2 mL elution buffer each (20 mM Tris-HCl pH 8, 25 mM reduced glutathione, 150 mM NaCl, 10 mM MgCl2). The relative IpaH amounts were quantified from a Coomassie-stained SDS-PAGE and the volumes subsequently used for the pull down assay adjusted accordingly to arrange for equal amount of GST-IpaH per sample. Between 8 and 24 μL of purified GST-IpaH protein was applied to 70 μL GST-Sepharose resin. This was used to pull down RNF213 from cell lysates. The cell lysates were prepared from RNF213 knockout MEFs, complemented with GFP-RNF213 (human). GFP-RNF213 expression was induced by the addition of 1 μM doxycycline 18 hours prior to harvesting the cells. The cells were washed in PBS and collected in lysis buffer (20 mM Tris-HCl pH 8, 150 mM NaCl, 0.1% Triton X-100, 5 mM EDTA, protease inhibitor tablet). The cell lysate was applied to the GST-IpaH on beads, the beads were washed three times, proteins were eluted from the resin in 100 μL elution buffer as above, and used for immunoblot analysis.

Pull down assays to test prospective binding mutants of RNF213 or IpaH1.4 were performed analogously, except for using bacterial lysates containing overexpressed GST-IpaH1.4 (wild type or binding mutants), rather than purified protein. For these experiments, 50 μL bacterial lysate was applied to 25 μL GST-Sepharose resin. Flag-GFP-RNF213 (wild type or binding mutants) or FLAG-GFP control containing cell lysates were prepared from 293ET cells transfected with the PiggyBac-RNF213 plasmid, including overnight incubation with doxycycline. Proteins were not eluted from the washed beads; instead, the bead suspension was boiled in Laemmli buffer and used for SDS-PAGE and immunoblot analysis.

### Mass spectrometry

Co-precipitation experiments to identify potential interactors of IpaH-family were performed similarly to the pull down assay as described above. GST beads (40 μL per specimen) were pre-charged with GST-IpaH proteins as described above. Then, 1×10^8^ HeLa cells were lysed and the lysates were added to each aliquot of IpaH-bound beads. Protein pull-downs were performed at 4°C with agitation for 2 hours. Beads were then washed at least 4 times in cold wash buffer (20 mM Tris-HCl, pH7.4, 150 mM NaCl, 0.1% Triton X-100, 5% glycerol, 5 mM EDTA, 1 mM PMSF, 1 mM benzamidine, 2 μg/mL aprotinin, 5 μg/mL leupeptin, 1 mM DTT). Buffer was exchanged to 100 mM ammonium bicarbonate and proteins were reduced in the presence of 10 mM DTT and alkylated with 55 mM iodoacetamide in a total volume of 30 μL. Proteins were then digested on-beads with 1 μg trypsin (Promega) for 18 hours at 37°C. Peptides were subsequently acidified by adding 4 μL of 2% formic acid. The sample was spun down at 14,000g for 5 min and 20 μL of supernatant was transferred into a vial. The peptide fractions (7 μL) were analysed by nano-scale capillary LC-MS/MS using an Ultimate U3000 HPLC (ThermoScientific Dionex) to deliver a flow of approximately 300 nL/min. A μ-precolumn cartridge C18 Acclaim PepMap 100 (5 μm, 300 μm x 5 mm) (ThermoScientific Dionex) trapped the peptides prior to separation on a C18 Acclaim PepMap100 (3 μm, 75 μm x 250 mm) (ThermoScientific Dionex, San Jose, USA). Peptides were eluted with a 90-minute gradient of acetonitrile (5% to 40%). The analytical column outlet was directly interfaced, via a modified nano-flow electrospray ionisation source, with a hybrid linear quadrupole ion trap mass spectrometer (Orbitrap LTQ Velos, ThermoScientific). Data analysis was carried out, using a resolution of 60,000 for the full MS spectrum, followed by ten MS/MS spectra in the linear ion trap. MS spectra were collected over an m/z range of 200–1800. MS/MS scans were collected using threshold energy of 35 eV for collision-induced dissociation. LC-MS/MS data were then searched against the Uniprot database using Mascot (Matrix Science). Database search parameters were set with a precursor tolerance of 10 ppm and a fragment ion mass tolerance of 0.8 Da. One missed enzyme cleavage was allowed and variable modifications for oxidized methionine, carbamidomethyl and phopho-(STY) were included. Two biological replicates of this experiment were performed.

### Specimen preparation for cryoEM

Recombinant RNF213 and IpaH1.4/2.5 were purified as described above. RNF213 was mixed with 5-fold molar excess of IpaH1.4 or IpaH2.5 and incubated on ice for 30 minutes. The final concentration of proteins was around 2 μM RNF213 and 10 μM IpaH1.4/2.5. All-gold grids (UltrAuFoil R 1.2/1.3, 300-mesh, QuantiFoil Micro Tools GmbH) (Russo & Passmore, 2014) were used for all specimens. The grids were plasma cleaned in an atmosphere of 9:1 Ar:O2 for 2 minutes at 70% power, 30 sccm gas flow in a Fischione 1070 plasma chamber. A humidified manual plunger of the Talmon type^60^, situated in a 4°C cold room was used to vitrify the specimens. Three microliters of each specimen were applied to the foil side of the grid and blotted from the same side for 10 seconds using Whatman No. 1 filter paper. The grid was then immediately plunged into liquid ethane, held at 93 Kelvin in a temperature-controlled cryostat.^61^ All grids were stored in liquid nitrogen until use.

### CryoEM data collection

Both datasets were collected at the electron Bio-Imaging Centre (eBIC) at Diamond Light Source, Harwell, UK. The dataset for the RNF213–IpaH1.4 complex was collected on a Titan Krios (ThermoFisher Scientific) electron cryomicroscope, equipped with a Schottky X-FEG operated at 300 keV and a Selectris X (ThermoFisher Scientific) imaging filter with a post-filter Falcon 4i (ThermoFisher Scientific) direct electron detector. The filter was used with a 10 eV slit and 4.56 second exposures, corresponding to 32 e^−^/Å^2^ fluence, were acquired in EER format at nominal magnification 130,000×, corresponding to 0.921 Å calibrated pixel size. The dataset for the RNF213–IpaH2.5 complex was collected on a Titan Krios electron cryomicroscope, equipped with a Schottky X-FEG operated at 300 keV and a BioQuantum (Gatan) imaging filter with a post-filter K3 (Gatan) direct electron detector. The filter was used with a 20 eV slit and 1.75 second exposures, corresponding to 40 e^−^/Å^2^ fluence, were acquired in TIFF format in super-resolution mode with 2-fold binning at nominal magnification 105,000×, corresponding to 0.825 Å calibrated pixel size. Both datasets were acquired using aberration-free image shift within a 6 μm range and correspond to 48 hours of data collection each. Data collection parameters are summarized in **Supplementary Table S1**.

### CryoEM data processing

Both datasets were processed using a combination of RELION-5.0^62^ and cryoSPARC-4.4^63^. The data processing strategy is summarized in **Extended Data Figure 7**. Briefly, all cryoEM movies were imported in RELION and motion corrected. The EER frames for the RNF213–IpaH1.4 dataset were grouped into 36 fractions, each corresponding to 0.9 e^−^/Å^2^ of irradiation. The contrast transfer functions were estimated using CTFFIND-4.1.^64^ Initial particle picking was performed using the Laplacian-of-Gaussian (LoG) picker in RELION with diameter range from 200 Å to 250 Å and minimal threshold at –2 standard deviations. The picked particles were extracted into 512-pixel boxes and binned by a factor of 4. The extracted particles were imported into cryoSPARC for 2D and 3D classification, where false picks and damaged particles were removed. The selected particles yielded an initial reconstruction of RNF213. This was low-pass filtered to 20 Å and used for template-based particle picking in RELION. The newly picked particles were extracted into 512-pixel boxes and binned by a factor of 2. The extracted particles were imported into cryoSPARC again for 2D and 3D classification. A selected subset of particles, that yielded a high-resolution reconstruction of RNF213, with extra low-resolution density corresponding to the IpaH1.4/2.5 LRR at partial occupancy were imported back into RELION and subjected to optical aberration refinement^65^ and Bayesian polishing^66^. The particles were then subjected to 3D refinement with regularization by the Blush algorithm.^67^ Masks encompassing the E3 domain of RNF213 and the IpaH1.4/2.5 LRR was created and used for masked 3D classification without alignment to remove particles without IpaH1.4/2.5 bound, followed by local 3D refinement with regularization by the Blush algorithm. This was repeated iteratively with progressively tighter masks, and the particles were re-sampled as necessary as resolution improved, until the final mask encompassed only IpaH1.4/2.5 and the RNF213 RING domain. All other RNF213 domains (ATPase, stalk, CBM) were also locally refined independently, as previously described.^14^ A composite map was produced from all refined domains, and this was used for model building. Graphics were generated in ChimeraX.^68^

### Atomic model building and refinement

The initial atomic models were created by docking the structures of human RNF213 (PDB 8S24)^14^ and of IpaH1.4/2.5 LRR (PDB 7YA7/7YA8)^69^ into the composite cryoEM maps (comprising the combination of the locally refined regions of the complex). The models were manually refined in ISOLDE^70^, followed by real-space refinement in PHENIX^71^ with secondary structure restraints and Ramachandran restraints.

## Acknowledgements

This work was supported by the Medical Research Council as part of United Kingdom Research and Innovation (U105170648), the Wellcome Trust (222503/Z/21/Z), the German Research Foundation (SCHA 2458/1-1) and a SNSF postdoc mobility grant to PMH (P400PB_191083). We acknowledge Diamond for access and support of the cryoEM facilities at eBIC, proposal BI31336, funded by the Wellcome Trust, MRC and BBSRC. We also thank the MRC LMB cryoEM, scientific computing, light microscopy, insect cell culture, mass spectrometry, and flow cytometry facilities for their support. We thank Johannes Schwab and Dari Kimanius for helpful advice on cryoEM data processing, Alison Lane for helpful advice on primer design, Christopher Tang, John Rohde, Neil Alto and David Holden for bacterial strains, and Michael Hensel for sharing plasmids.

## Data availability

The cryoEM maps and the refined atomic models have been deposited in Electron Microscopy Data Bank and the Protein Data Bank under the accession codes EMD 50913–50929 and PDB 9G08 and 9G09, respectively. Other data is available within the article and its associated supplementary information files.

## Author contributions

KN, KBB, CP, PP, ACC, FS, PMH and EGO performed experiments and analyzed the results. KN, KB and FR wrote the manuscript with input from all authors.

## Extended Data Figure legends

**Extended Data Figure 1.**
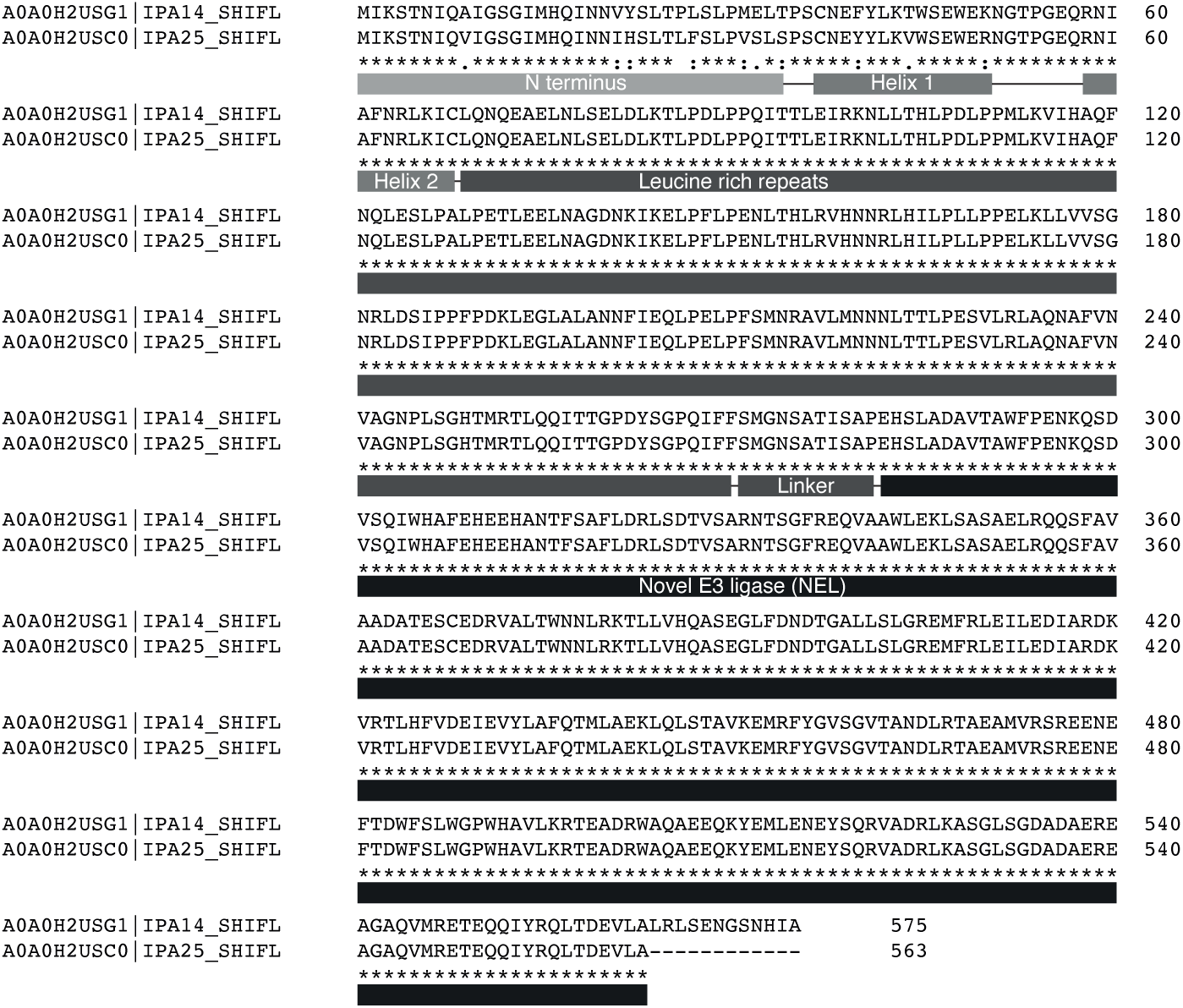
Sequence alignment of *Shigella flexneri* IpaH1.4/2.5. The IpaH1.4 and IpaH2.5 protein sequences from *S. flexneri*, as listed in UNIPROT, were aligned using Clustal Omega. The structural domains are annotated below the sequences. IpaH1.4 and IpaH2.5 only differ in (1) the unstructured N-terminus, (2) the first alpha-helix preceding the leucine rich repeat, and (3) the C-terminal extension following the NEL domain. The leucine rich repeat, the NEL domain, and the interconnecting linker are 100% identical between the two proteins.

**Extended Data Figure 2.**
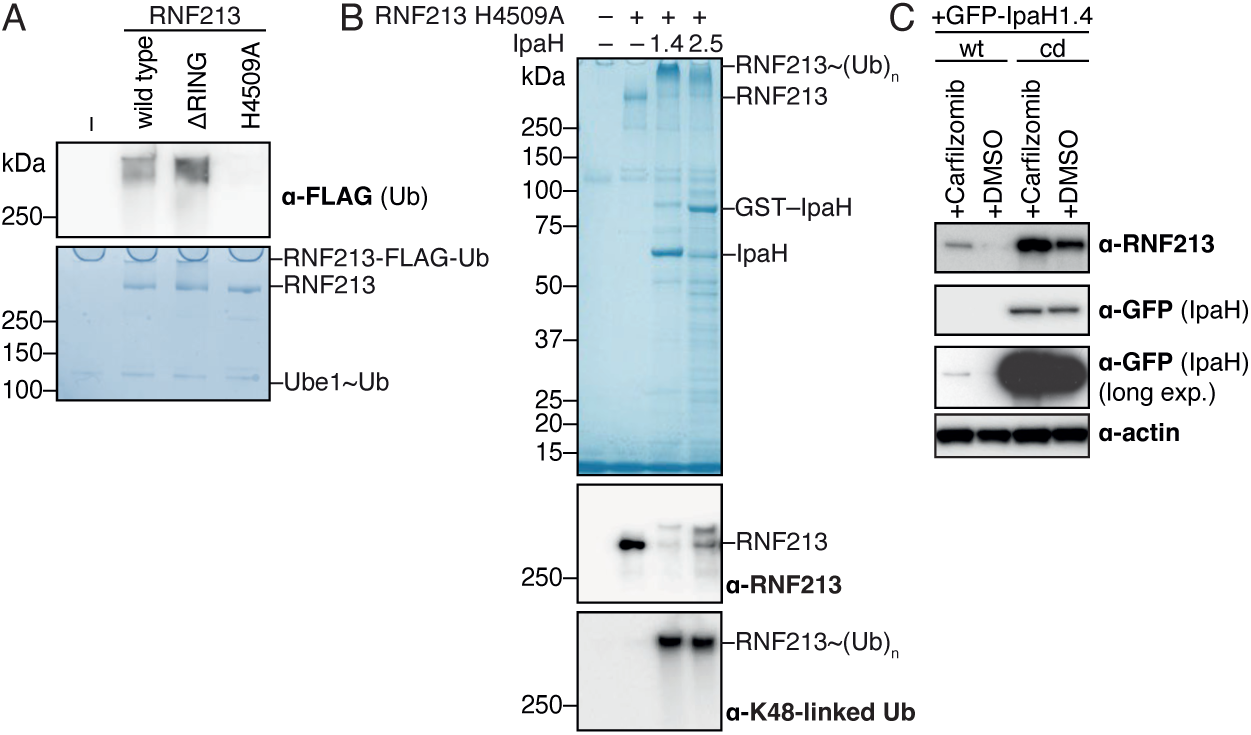
Choice of RNF213 mutant, lacking auto-ubiquitylation activity, for IpaH1.4/2.5-mediated ubiquitylation experiments. (A) Auto-ubiquitylation assays of the indicated RNF213 variants incubated with Flag-tagged ubiquitin, UBE1 and UBCH5C (as described in the Methods). Upper: anti-Flag immunoblot. Lower: Coomassie stained SDS-PAGE. (B) *In vitro* ubiquitylation reaction containing Flag-tagged ubiquitin, UBE1, UBCH5C, the indicated IpaH proteins and enzymatically inactive RNF213 H4509A as substrate. Upper panel, Coomassie stained SDS-PAGE. Middle panel, immunoblot for RNF213. Bottom panel, immunoblot for K48-linked ubiquitin. (C) Immunoblot analysis of HeLa cell lysates expressing the indicated GFP-tagged IpaH proteins after treatment with proteasome inhibitor (Carfilzomib) or control (DMSO). Actin, loading control; cd, catalytically dead (E3 inactive) variant; wt, wild-type.

**Extended Data Figure 3.**
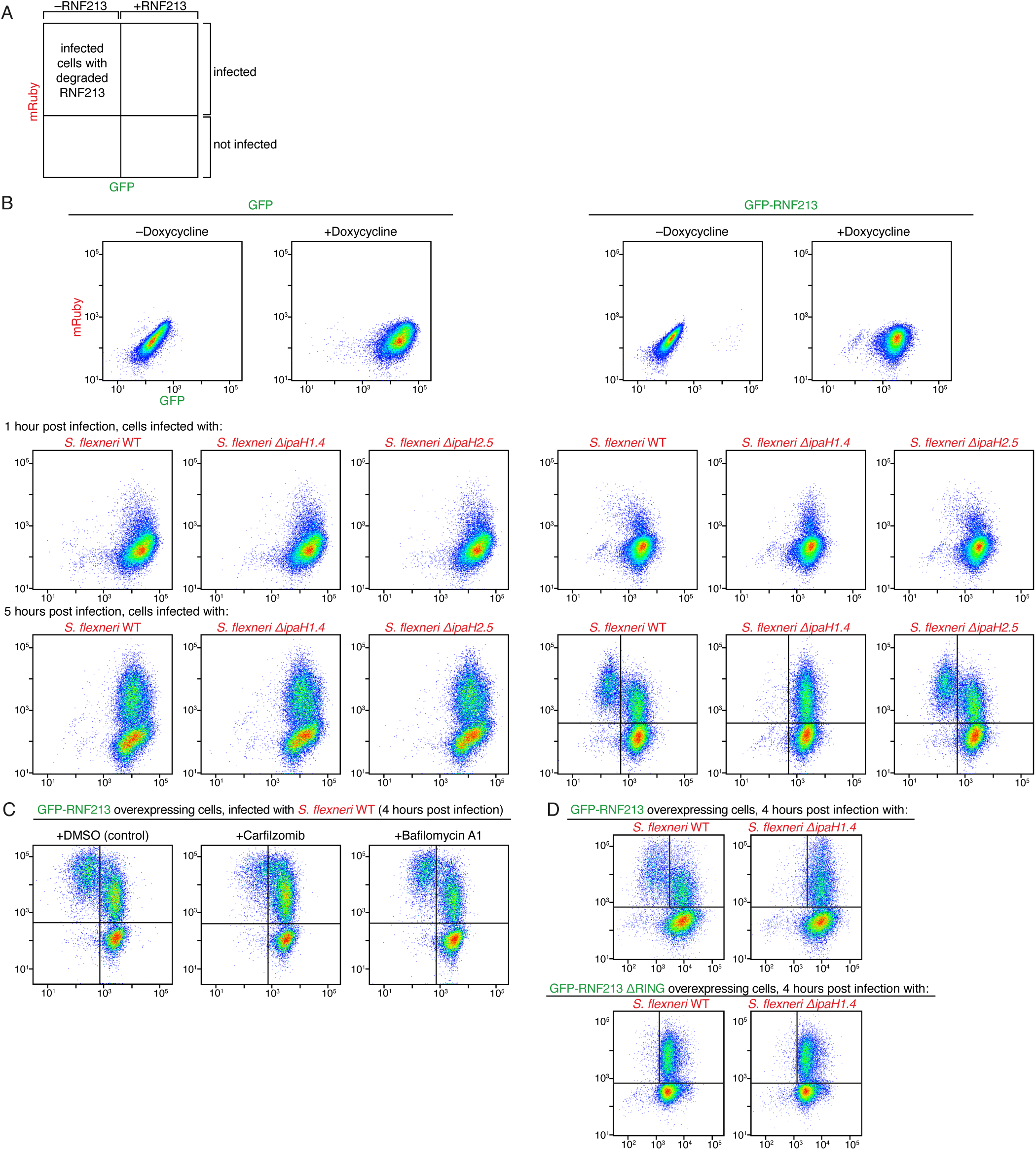
Quantification of *S. flexneri* induced degradation of RNF213 by flow cytometry. (A) Cartoon representation of flow cytometry scatter plots below, indicating the identity of events observed in each quadrant. (B) Flow cytometry of GFP (left) and GFP-RNF213 (right) expressing MEFs induced with or without Doxycycline as indicated (top row); after induction with Doxycycline at 1h (middle row) or 5h post infection (bottom row) with the indicated strains of Ruby-labelled *S. flexneri*. (C) Flow cytometry of GFP-RNF213 expressing MEFs induced with Doxycycline and treated with the indicated inhibitors at 4h post infection with Ruby-labelled *S. flexneri*. (D) Flow cytometry of GFP-RNF213 (upper) and GFP-RNF213 ΔRING (lower) expressing MEFs at 4 hours post infection with the indicated strains of *S. flexneri.* All plots are on the same scale and include 30,000 events/plot.

**Extended Data Figure 4.**
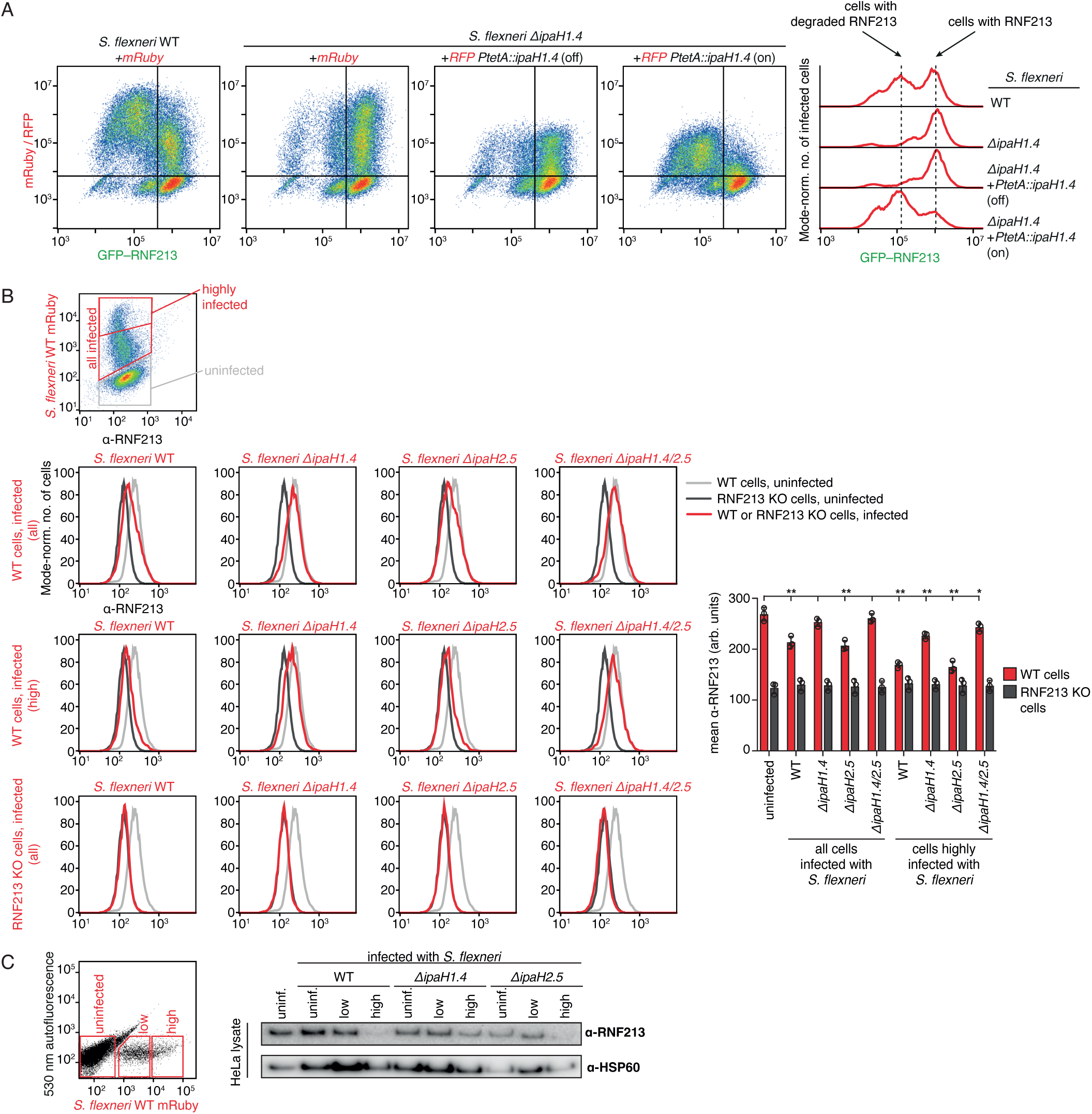
*Shigella flexneri* uses IpaH1.4 to degrade RNF213. (A) Complementation of *S. flexneri ΔipaH1.4* with plasmid-encoded *IpaH1.4* rescues degradation of RNF213 in infected host cells. Left: Flow cytometry of MEFs expressing GFP-RNF213 under control of a Doxycycline-inducible promoter, induced with Doxycycline, and infected with the indicated strains of *S. flexneri* harboring plasmids encoding constitutively expressed mRuby or TagRFP-T and anhydrotetracycline-inducible *IpaH1.4*, induced with anhydrotetracycline (ATc) as indicated (off – not induced, on – induced). Cells were analyzed by flow cytometry at 5 hours post infection. Right: histogram plots of GFP-RNF213 intensity in infected cells. (B) Analysis of endogenous RNF213 levels in *S. flexneri*-infected HeLa cells by flow cytometry. Wild type or RNF213 KO HeLa cells infected with the indicated strains of *S. flexneri* expressing mRuby were stained for endogenous RNF213. Levels of RNF213 are depicted as histograms from the indicated cell populations. The difference in staining between RNF213 WT and KO cells is indicative of the specific contribution of RNF213 to the assay. Infection with either *S. flexneri* WT or *ΔIpaH2.5*, but not *ΔIpaH1.4*, causes loss of RNF213 protein. (C) Upper: Gating strategy for fluorescence-activated cell sorting (FACS) of HeLa cells. HeLa cells were infected with the indicated mRuby-expressing *S. flexneri* strains. At 4 hours post infection the three indicated populations were isolated by FACS. Lower: Immunoblot analysis of RNF213 in the indicated cell populations. HSP60 – loading control.

**Extended Data Figure 5.**
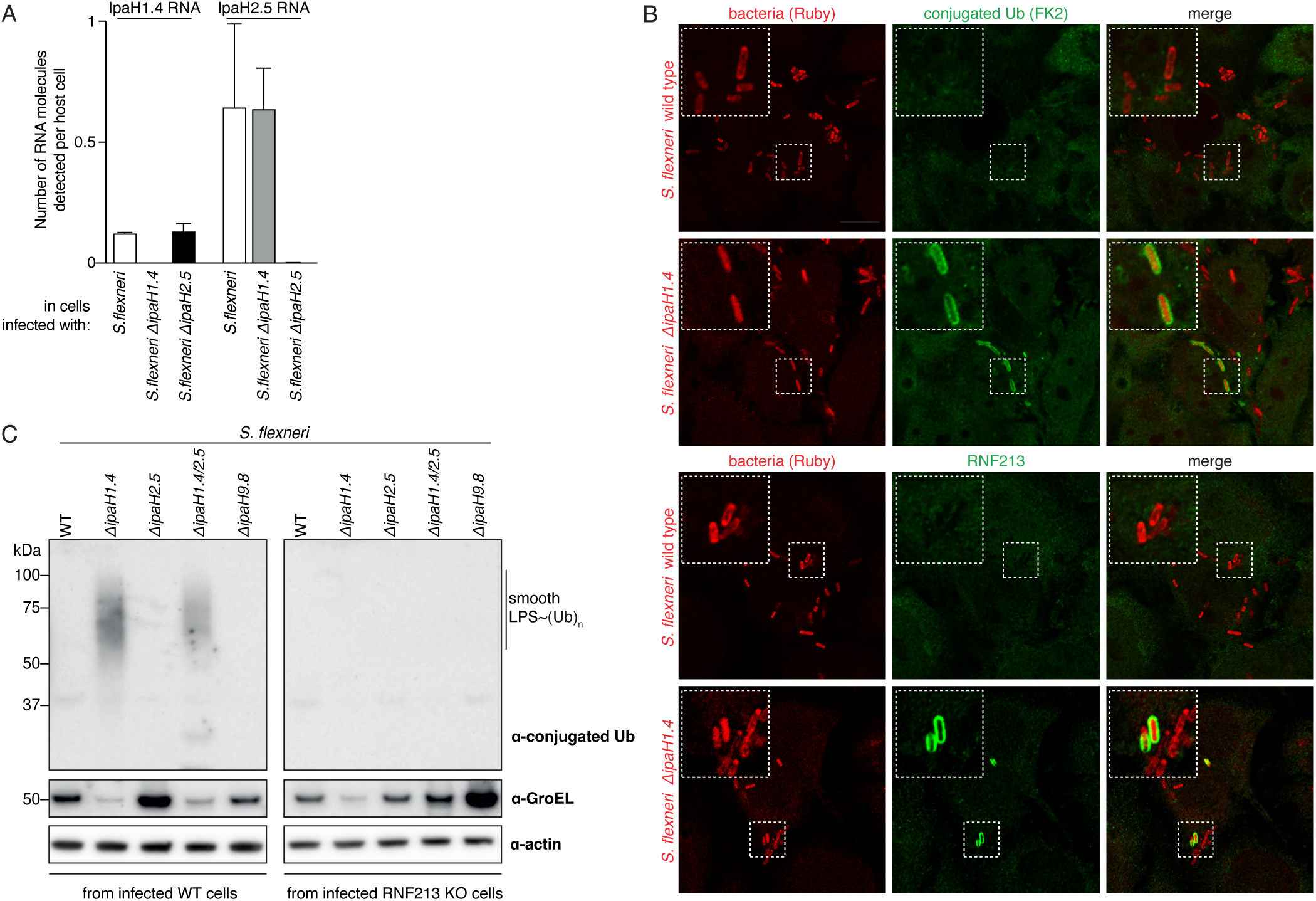
IpaH1.4 antagonizes RNF213-mediated LPS ubiquitylation in mouse and human cells. (A) Detection of IpaH1.4 and IpaH2.5 transcripts in *Shigella flexneri*. The bars show mean values, and the error bars denote standard deviation between *n*=2 replicate experiments. The number of RNA molecules per host cell is estimated by dividing the number of RNA molecules (calculated relative to a plasmid for normalization, as described in the Methods) by the total number of cells used for each measurement (∼10^5^). This represents a lower bound for the actual number of transcripts per bacterium due to the limited efficiency of the RNA extraction method. (B) Expanded view of Figure 3B. Representative confocal micrographs of HeLa cells at 4 hours post infection with the indicated strains of Ruby-labeled *Shigella flexneri*, immunostained for conjugated ubiquitin (FK2) (left), or RNF213 (right). Dashed squares are magnified 2.5× in the insets. Scale bar, 10 μm. (C) Immunoblot analysis of the indicated strains of *S. flexneri* extracted from wild-type MEFs (left, same as Figure 3D) and RNF213 KO MEFs (right) at 4 h post-infection. The LPS fractions were isolated by heat clearance of the bacterial lysates and probed for conjugated ubiquitin with FK2 antibody. The loading control (GroEL) was obtained from non-heat cleared bacterial lysates. The host cell loading control (actin) was obtained from clarified host cell lysates.

**Extended Data Figure 6.**
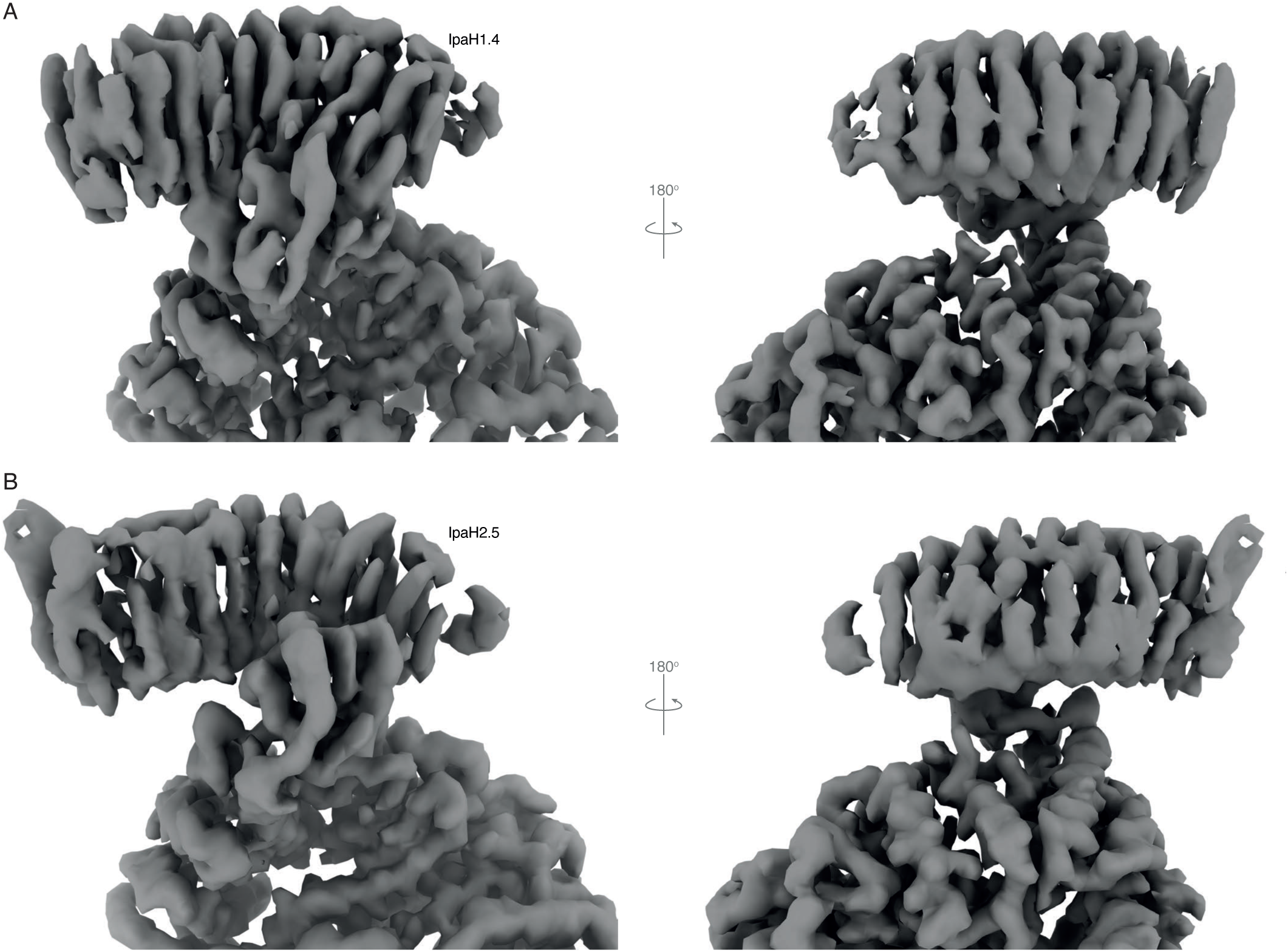
Structure of the IpaH2.5–RNF213 complex. Comparison between the cryoEM maps of (A) IpaH1.4–RNF213 and (B) IpaH2.5–RNF213 complexes.

**Extended Data Figure 7.**
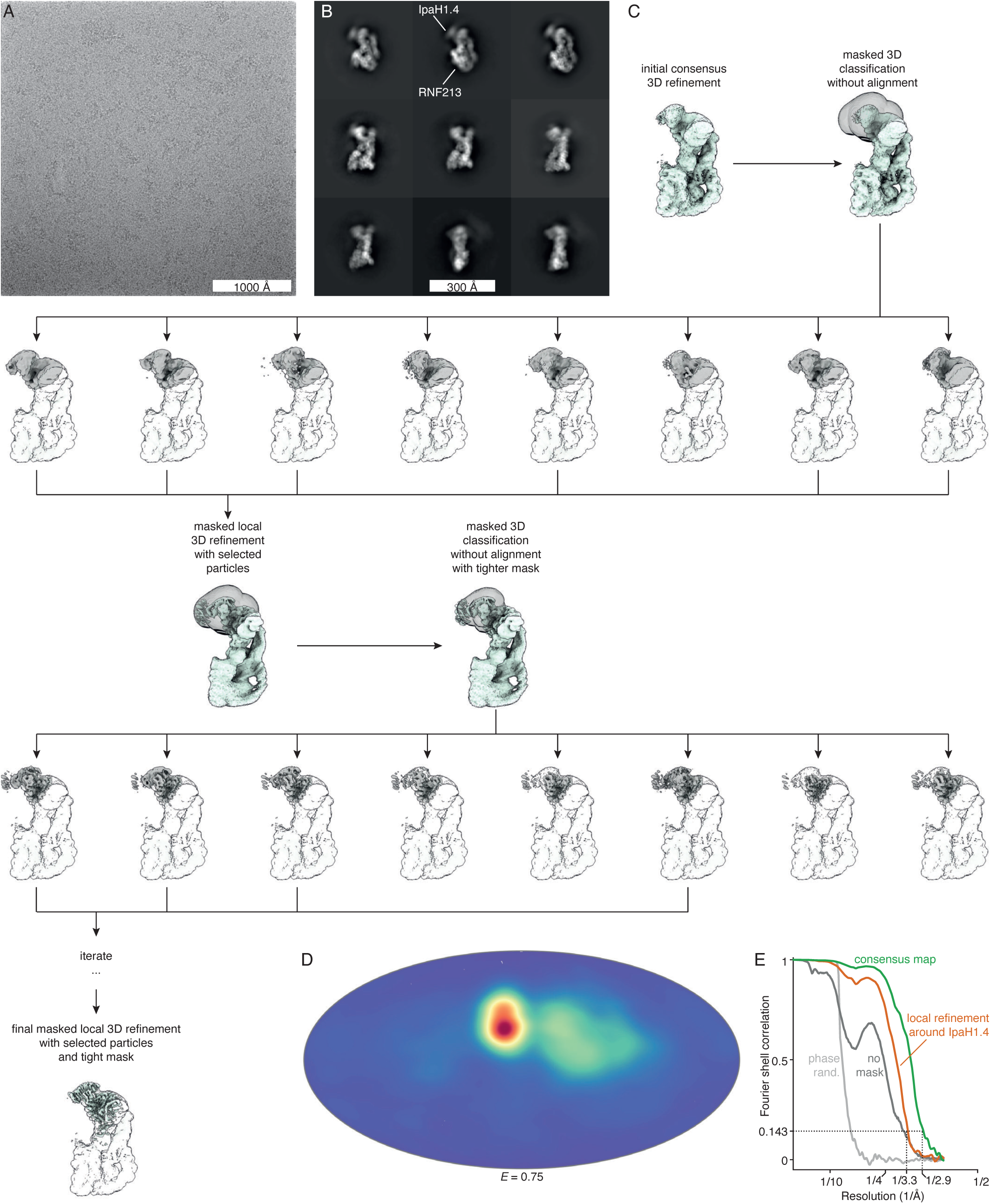
CryoEM data processing. (A) A representative micrograph, acquired at 1.5 μm defocus, from the IpaH1.4–RNF213 dataset, low-pass filtered to 10 Å for display. (B) Representative 2D class averages of the IpaH1.4–RNF213 complex. (C) The flow chart summarizes the three-dimensional processing strategy, aimed at removing RNF213 particles without IpaH1.4 bound and at improving the resolution around the flexible region of the IpaH1.4 LRR and RNF213 RING. All processing steps after the initial consensus 3D refinement were performed in RELION. (D) Orientation distribution of the particles used in the final reconstruction of the IpaH1.4–RNF213 complex, plotted on a Mollweide projection. The efficiency of the orientation distribution, *E*, calculated using cryoEF^75^, is 0.75. (E) Fourier shell correlation as function of resolution is plotted for the best consensus masked map (green), the best masked map for the locally refined region of the IpaH1.4 LRR and RNF213 RING (orange), the unmasked consensus map (dark grey), and the phase-randomized consensus map (light grey).

**Extended Data Figure 8.**
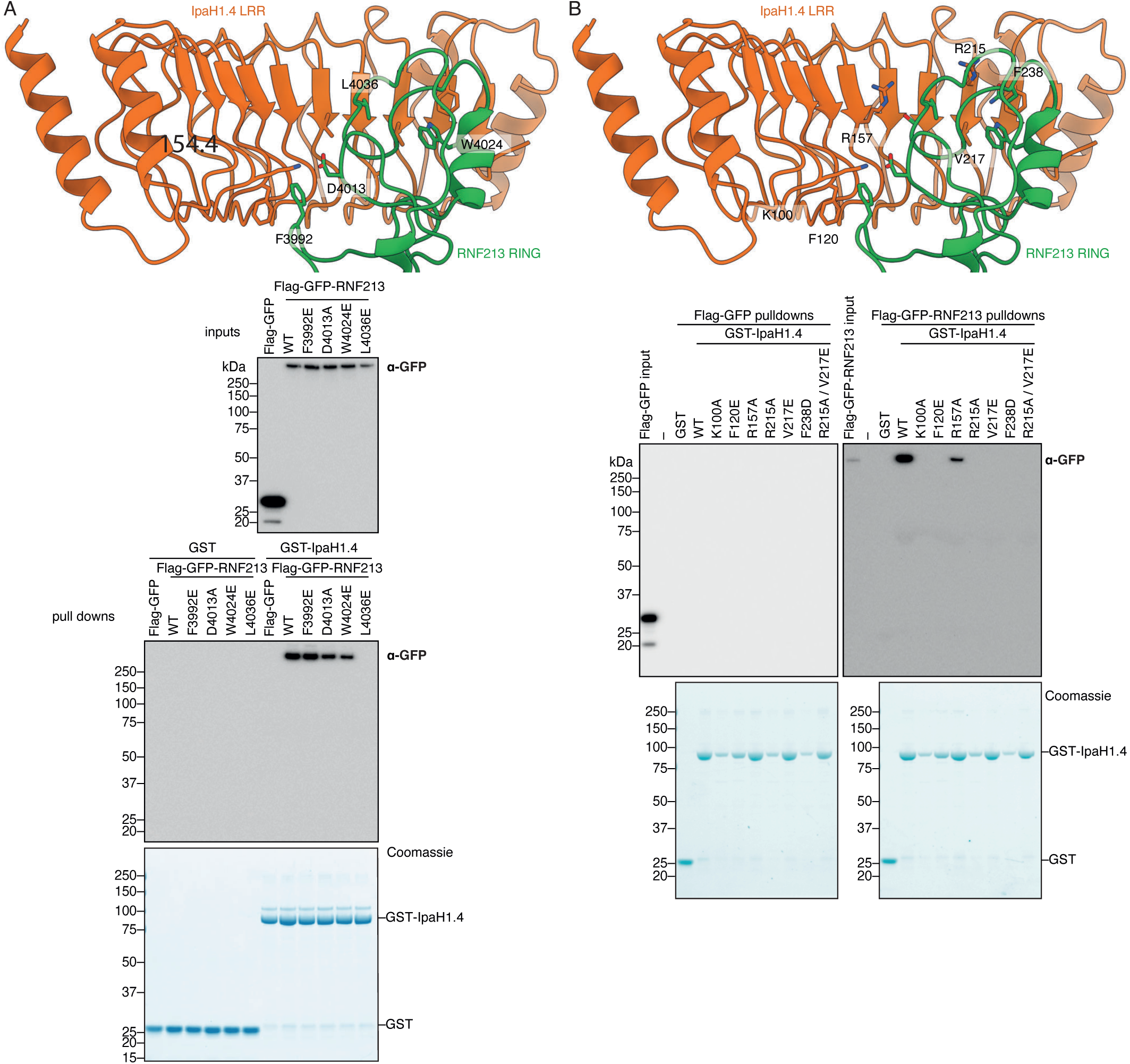
Validation of the IpaH1.4–RNF213 binding interface. (A) Pulldown of the indicated Flag-GFP-tagged RNF213 mutant proteins from cell lysates upon incubation with purified GST-IpaH1.4. Mutated residues and their interaction partners are highlighted in the cartoon. (B) Pulldown of Flag-GFP-RNF213 (or Flag-GFP as negative control) from cell lysates upon incubation with the indicated GST-IpaH1.4 mutant proteins (or GST as negative control). Mutated residues and their interaction partners are highlighted in the cartoon.

## Supplementary Figures

**Supplementary Figure 1.**
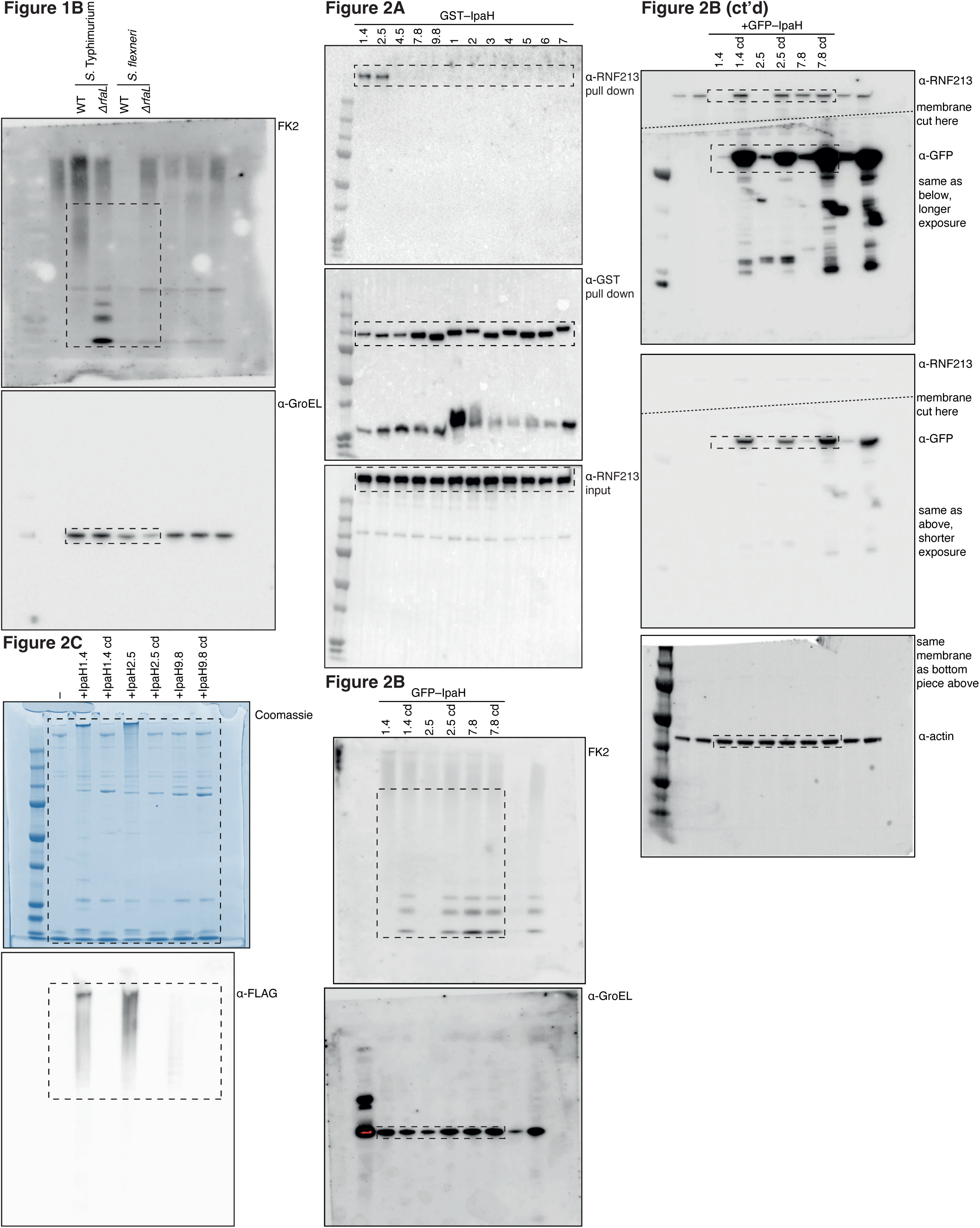
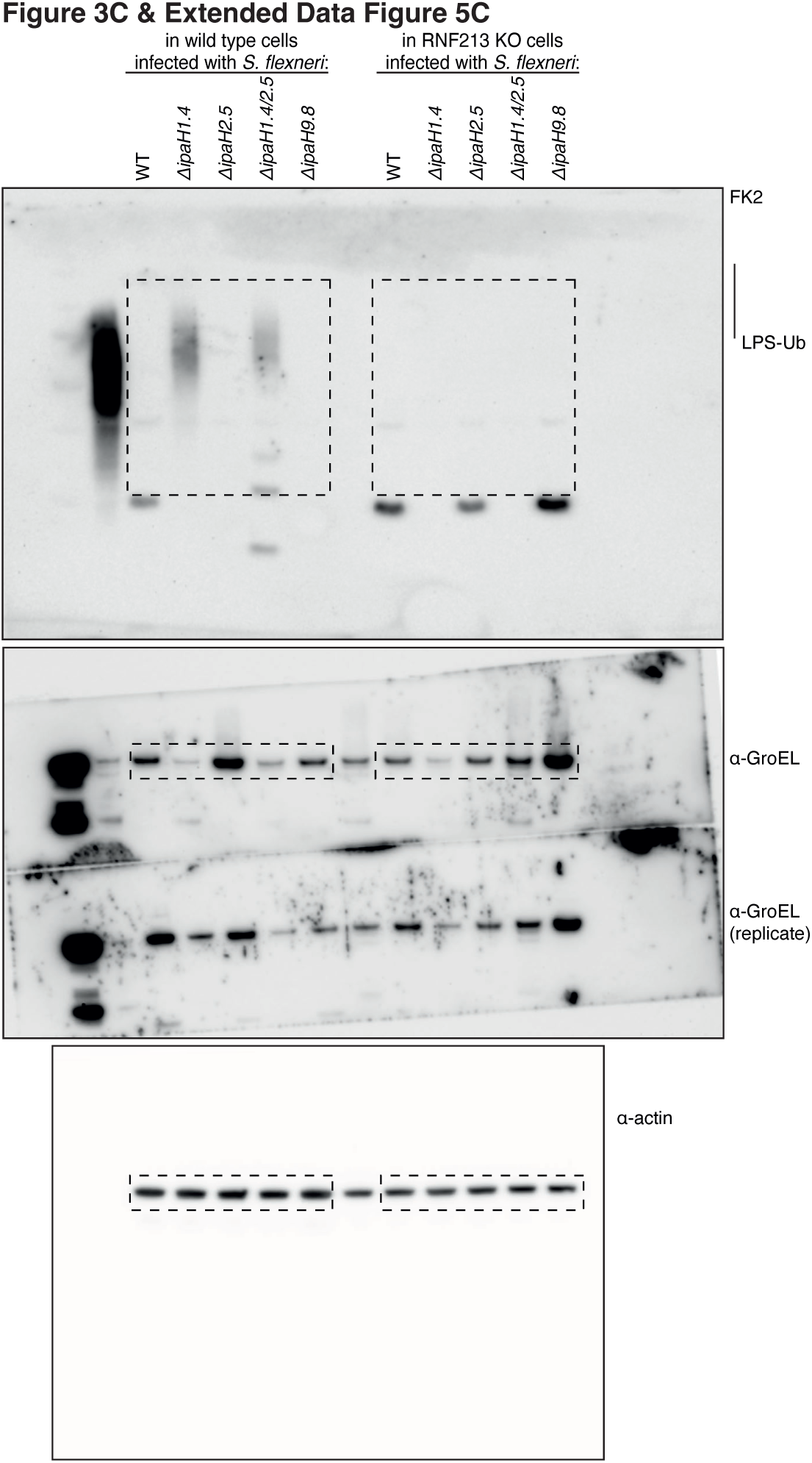
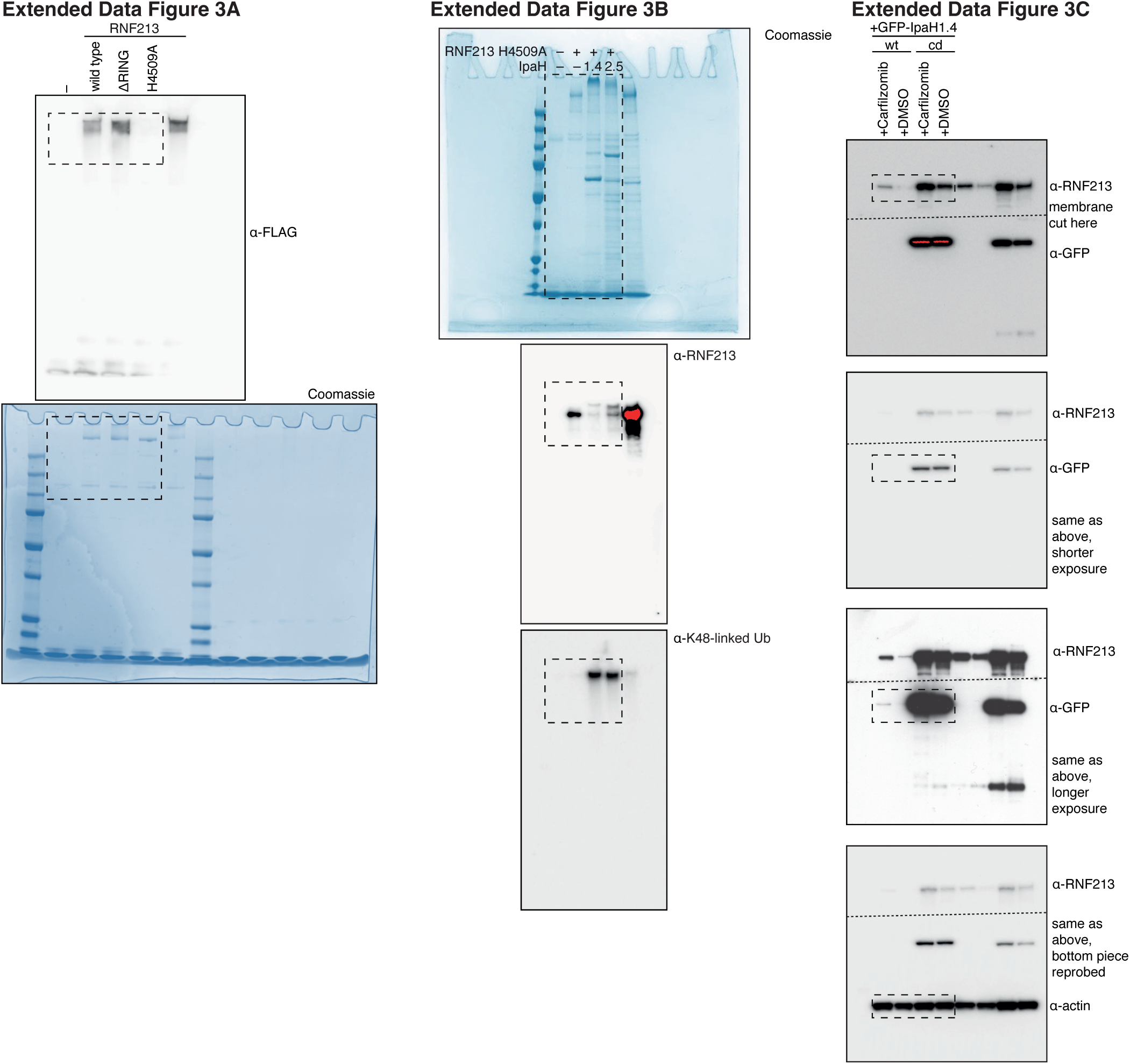
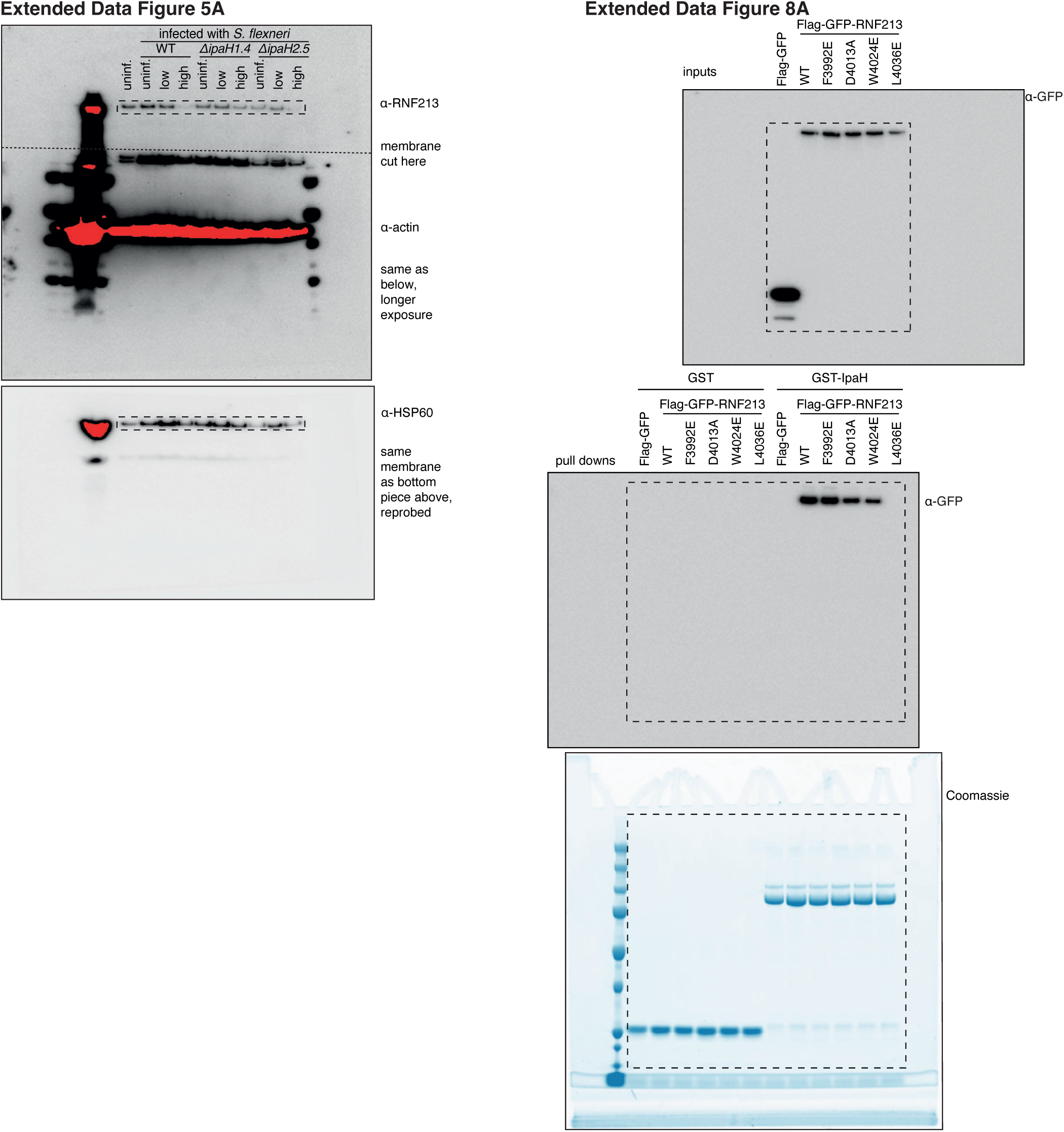
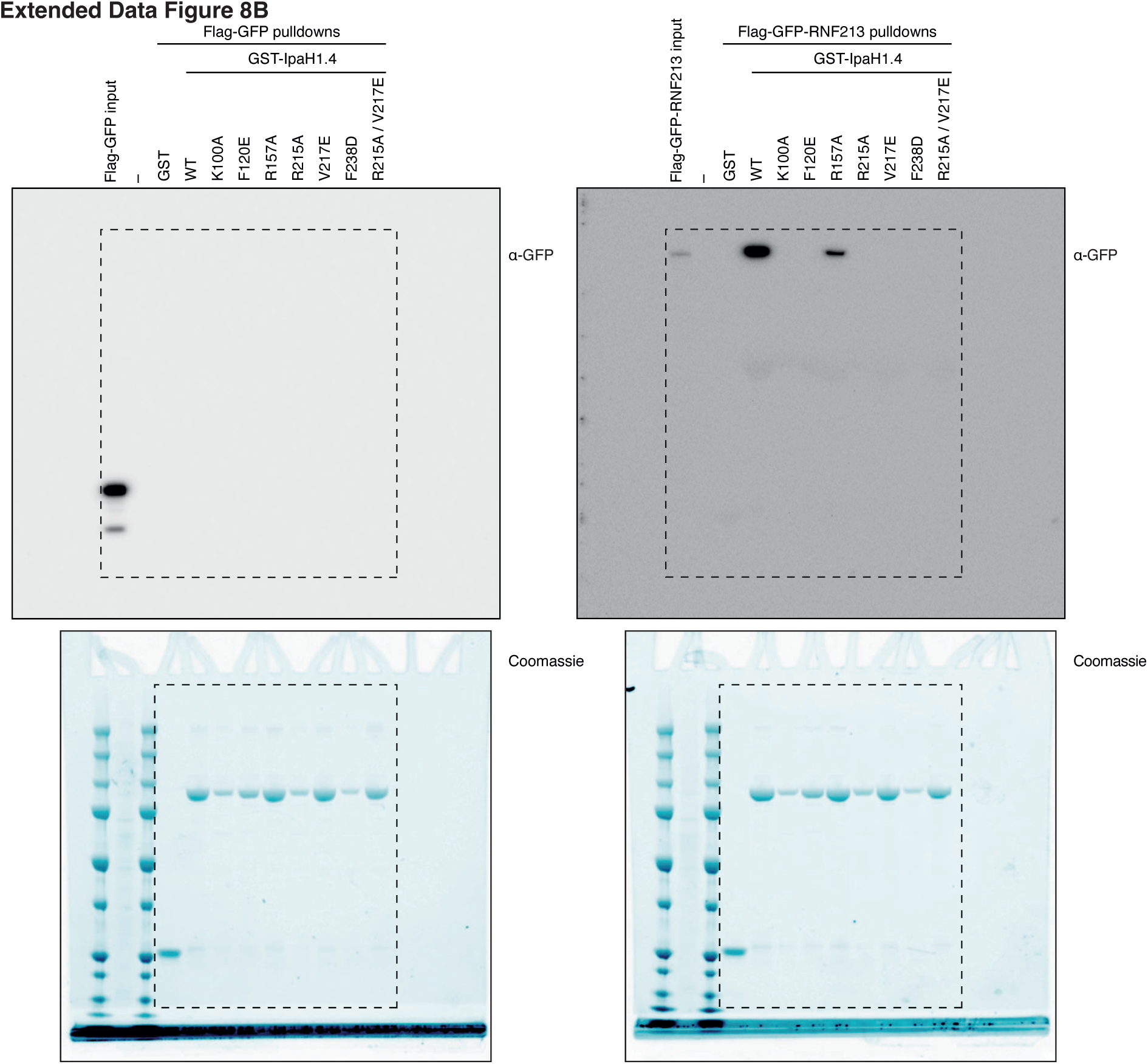
Source data for cropped immunoblots and gels. Original images of the cropped immunoblots and gels shown in **Figures 1–3 and Extended Data Figures 2, 4, 5, 8**.

## Supplementary Tables

**Table S1.**
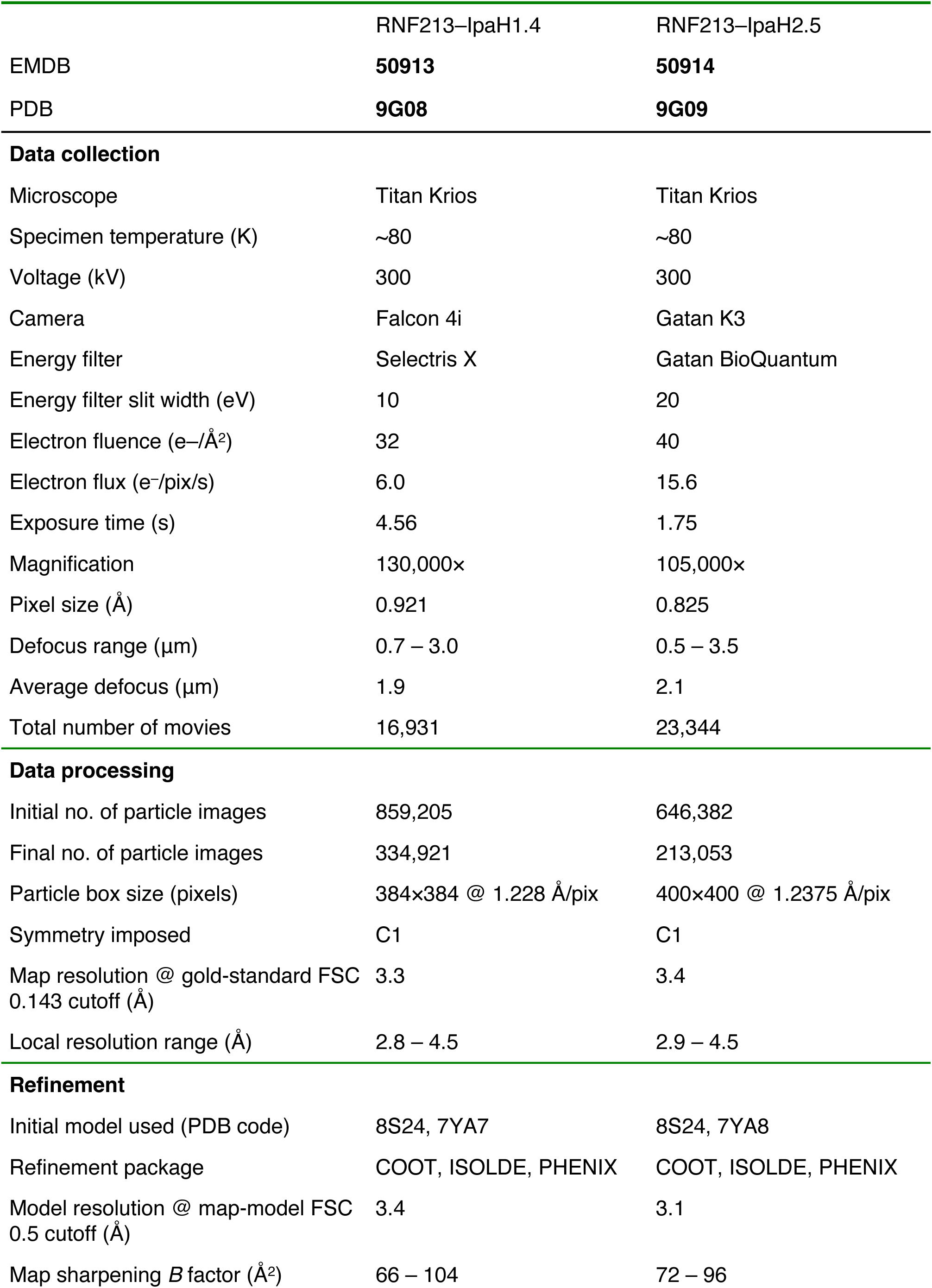
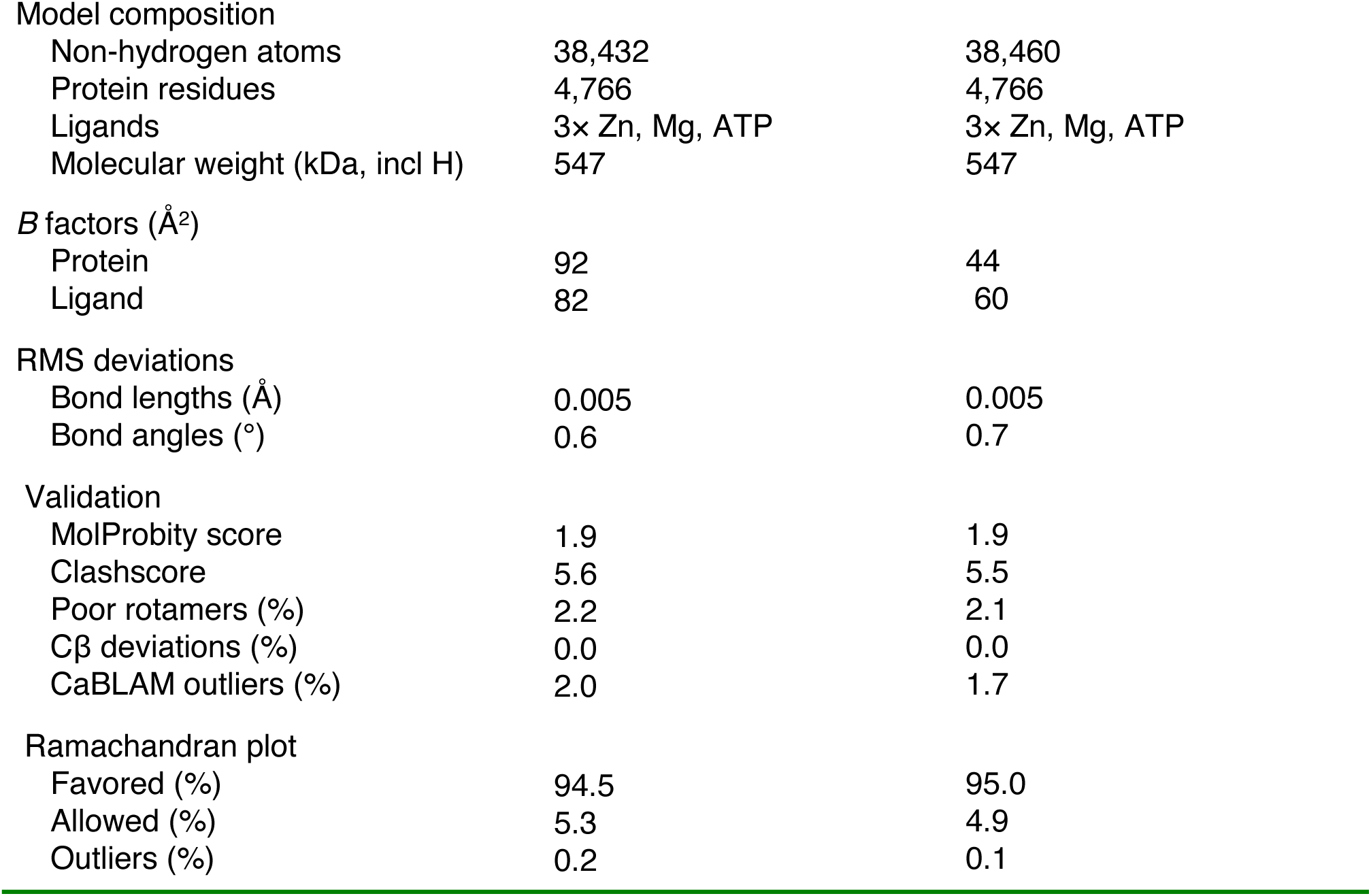
CryoEM data collection and processing, model refinement and validation statistics.

|  | RNF213–IpaH1.4 | RNF213–IpaH2.5 |
| --- | --- | --- |
| EMDB | **50913** | **50914** |
| PDB | **9G08** | **9G09** |
| **Data collection** |  |  |
| Microscope | Titan Krios | Titan Krios |
| Specimen temperature (K) | ~80 | ~80 |
| Voltage (kV) | 300 | 300 |
| Camera | Falcon 4i | Gatan K3 |
| Energy filter | Selectris X | Gatan BioQuantum |
| Energy filter slit width (eV) | 10 | 20 |
| Electron fluence ($e^-/\text{\AA}^2$) | 32 | 40 |
| Electron flux ($e^-/\text{pix/s}$) | 6.0 | 15.6 |
| Exposure time (s) | 4.56 | 1.75 |
| Magnification | 130,000× | 105,000× |
| Pixel size ($\text{\AA}$) | 0.921 | 0.825 |
| Defocus range ($\mu\text{m}$) | 0.7 – 3.0 | 0.5 – 3.5 |
| Average defocus ($\mu\text{m}$) | 1.9 | 2.1 |
| Total number of movies | 16,931 | 23,344 |
| **Data processing** |  |  |
| Initial no. of particle images | 859,205 | 646,382 |
| Final no. of particle images | 334,921 | 213,053 |
| Particle box size (pixels) | 384×384 @ 1.228 $\text{\AA}/\text{pix}$ | 400×400 @ 1.2375 $\text{\AA}/\text{pix}$ |
| Symmetry imposed | C1 | C1 |
| Map resolution @ gold-standard FSC 0.143 cutoff ($\text{\AA}$) | 3.3 | 3.4 |
| Local resolution range ($\text{\AA}$) | 2.8 – 4.5 | 2.9 – 4.5 |
| **Refinement** |  |  |
| Initial model used (PDB code) | 8S24, 7YA7 | 8S24, 7YA8 |
| Refinement package | COOT, ISOLDE, PHENIX | COOT, ISOLDE, PHENIX |
| Model resolution @ map-model FSC 0.5 cutoff ($\text{\AA}$) | 3.4 | 3.1 |
| Map sharpening $B$ factor ($\text{\AA}^2$) | 66 – 104 | 72 – 96 |
| Model composition |  |  |
| Non-hydrogen atoms | 38,432 | 38,460 |
| Protein residues | 4,766 | 4,766 |
| Ligands | 3× Zn, Mg, ATP | 3× Zn, Mg, ATP |
| Molecular weight (kDa, incl H) | 547 | 547 |
| *B* factors (Å<sup>2</sup>) |  |  |
| Protein | 92 | 44 |
| Ligand | 82 | 60 |
| RMS deviations |  |  |
| Bond lengths (Å) | 0.005 | 0.005 |
| Bond angles (°) | 0.6 | 0.7 |
| Validation |  |  |
| MolProbity score | 1.9 | 1.9 |
| Clashscore | 5.6 | 5.5 |
| Poor rotamers (%) | 2.2 | 2.1 |
| Cβ deviations (%) | 0.0 | 0.0 |
| CaBLAM outliers (%) | 2.0 | 1.7 |
| Ramachandran plot |  |  |
| Favored (%) | 94.5 | 95.0 |
| Allowed (%) | 5.3 | 4.9 |
| Outliers (%) | 0.2 | 0.1 |

**Table S2.**
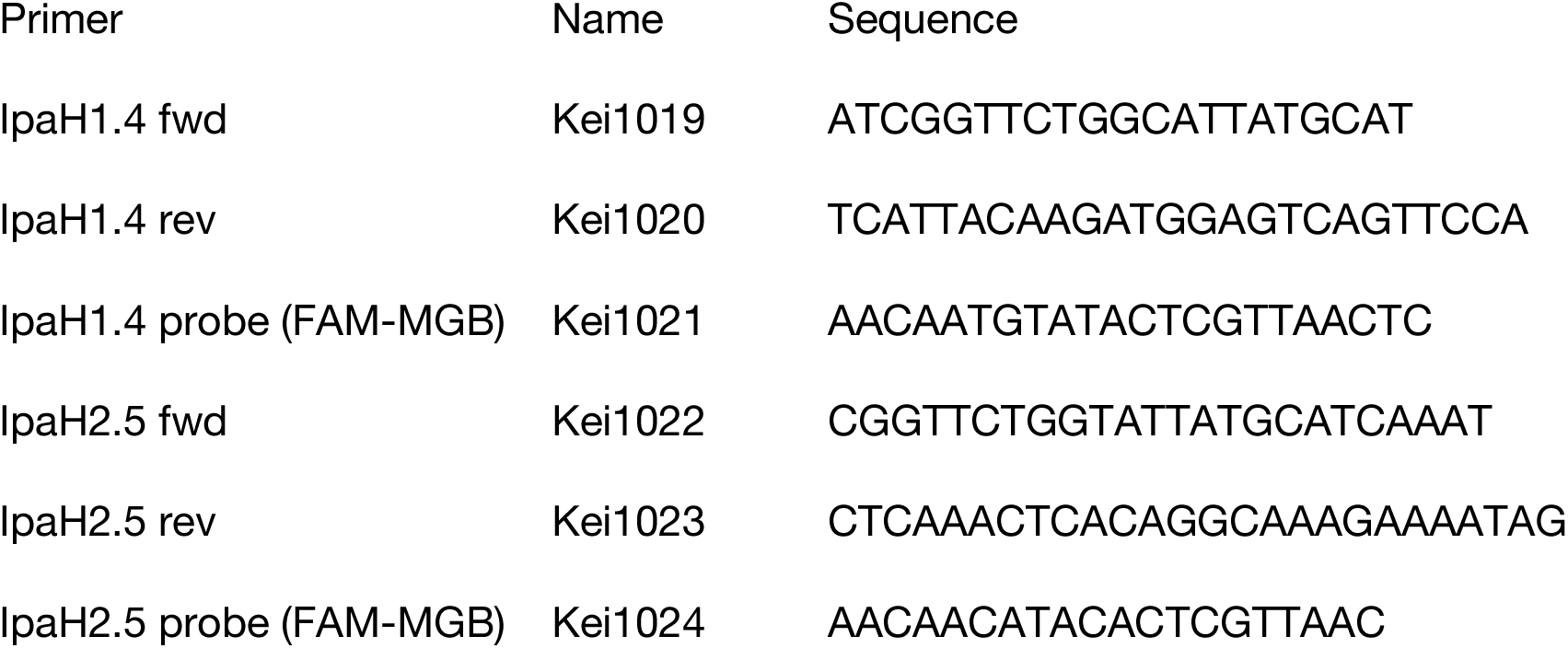
Primers used for detection of IpaH1.4/2.5 transcripts in infected cells.

**Table S3. Mass spectrometry-based identification of putative targets of IpaH1.4/2.5 from HeLa cell lysates**. The list of identified interaction partners is provided as a separate file.

## References

1. Casadevall, A. Evolution of Intracellular Pathogens. Annu. Rev. Microbiol. 62, 19–33 (2008).

2. Randow, F., MacMicking, J. D. & James, L. C. Cellular Self-Defense: How Cell-Autonomous Immunity Protects Against Pathogens. Science 340, 701–706 (2013).

3. Huang, J. & Brumell, J. H. Bacteria–autophagy interplay: a battle for survival. Nat Rev Microbiol 12, 101–114 (2014).

4. Thurston, T. L. M., Wandel, M. P., Muhlinen, N. von, Foeglein, Á. & Randow, F. Galectin 8 targets damaged vesicles for autophagy to defend cells against bacterial invasion. Nature 482, 414–418 (2012).

5. Perrin, A. J., Jiang, X., Birmingham, C. L., So, N. S. Y. & Brumell, J. H. Recognition of Bacteria in the Cytosol of Mammalian Cells by the Ubiquitin System. Curr Biol 14, 806–811 (2004).

6. Thurston, T. L. M., Ryzhakov, G., Bloor, S., Muhlinen, N. von & Randow, F. The TBK1 adaptor and autophagy receptor NDP52 restricts the proliferation of ubiquitin-coated bacteria. Nat Immunol 10, 1215–1221 (2009).

7. Wild, P. et al. Phosphorylation of the Autophagy Receptor Optineurin Restricts Salmonella Growth. Science 333, 228–233 (2011).

8. Zheng, Y. T. et al. The Adaptor Protein p62/SQSTM1 Targets Invading Bacteria to the Autophagy Pathway. J Immunol 183, 5909–5916 (2009).

9. Tumbarello, D. A. et al. Autophagy receptors link myosin VI to autophagosomes to mediate Tom1-dependent autophagosome maturation and fusion with the lysosome. Nat Cell Biol 14, 1024– 1035 (2012).

10. Otten, E. G. et al. Ubiquitylation of lipopolysaccharide by RNF213 during bacterial infection. Nature 594, 111–116 (2021).

11. Noad, J. et al. LUBAC-synthesized linear ubiquitin chains restrict cytosol-invading bacteria by activating autophagy and NF-κB. Nat Microbiol 2, 17063 (2017).

12. Wijk, S. J. L. van et al. Linear ubiquitination of cytosolic Salmonella Typhimurium activates NF-κB and restricts bacterial proliferation. Nat Microbiol 2, 17066 (2017).

13. Thery, F. et al. Ring finger protein 213 assembles into a sensor for ISGylated proteins with antimicrobial activity. Nat Commun 12, 5772 (2021).

14. Crespillo Casado, A., et al. Recognition of phylogenetically diverse pathogens through enzymatically amplified recruitment of RNF213. EMBO Rep. in press (2024).

15. Hernandez, D., Walsh, S., Sanchez, L. S., Dickinson, M. S. & Coers, J. Interferon-Inducible E3 Ligase RNF213 Facilitates Host-Protective Linear and K63-Linked Ubiquitylation of Toxoplasma gondii Parasitophorous Vacuoles. Mbio e01888–22 (2022) doi:10.1128/mbio.01888-22.

16. Walsh, S. C. et al. The bacterial effector GarD shields Chlamydia trachomatis inclusions from RNF213-mediated ubiquitylation and destruction. Cell Host Microbe (2022) doi:10.1016/j.chom.2022.08.008.

17. Matta, S. K. et al. Genome-wide and targeted CRISPR screens identify RNF213 as a mediator of interferon gamma–dependent pathogen restriction in human cells. Proc. Natl. Acad. Sci. 121, e2315865120 (2023).

18. Houzelstein, D. et al. The ring finger protein 213 gene (Rnf213) contributes to Rift Valley fever resistance in mice. Mamm Genome 1–8 (2021) doi:10.1007/s00335-020-09856-y.

19. Liu, W. et al. Identification of RNF213 as a Susceptibility Gene for Moyamoya Disease and Its Possible Role in Vascular Development. Plos One 6, e22542 (2011).

20. Kamada, F. et al. A genome-wide association study identifies RNF213 as the first Moyamoya disease gene. J Hum Genet 56, 34–40 (2011).

21. Ihara, M. et al. Moyamoya disease: diagnosis and interventions. Lancet Neurology 21, 747–758 (2022).

22. Baxt, L. A., Garza-Mayers, A. C. & Goldberg, M. B. Bacterial Subversion of Host Innate Immune Pathways. Science 340, 697–701 (2013).

23. Ogawa, M. et al. Escape of Intracellular Shigella from Autophagy. Science 307, 727–731 (2005).

24. Baxt, L. A. & Goldberg, M. B. Host and Bacterial Proteins That Repress Recruitment of LC3 to Shigella Early during Infection. Plos One 9, e94653 (2014).

25. Ashida, H. & Sasakawa, C. Shigella IpaH Family Effectors as a Versatile Model for Studying Pathogenic Bacteria. Front Cell Infect Mi 5, 100 (2015).

26. Rohde, J. R., Breitkreutz, A., Chenal, A., Sansonetti, P. J. & Parsot, C. Type III Secretion Effectors of the IpaH Family Are E3 Ubiquitin Ligases. Cell Host Microbe 1, 77–83 (2007).

27. Singer, A. U. et al. Structure of the Shigella T3SS effector IpaH defines a new class of E3 ubiquitin ligases. Nat Struct Mol Biol 15, 1293–1301 (2008).

28. Zhu, Y. et al. Structure of a Shigella effector reveals a new class of ubiquitin ligases. Nat Struct Mol Biol 15, 1302–1308 (2008).

29. Quezada, C. M., Hicks, S. W., Galan, J. E. & Stebbins, C. E. A family of Salmonella virulence factors functions as a distinct class of autoregulated E3 ubiquitin ligases. Proc National Acad Sci 106, 4864–4869 (2009).

30. Chou, Y.-C., Keszei, A. F. A., Rohde, J. R., Tyers, M. & Sicheri, F. Conserved Structural Mechanisms for Autoinhibition in IpaH Ubiquitin Ligases. J Biol Chem 287, 268–275 (2012).

31. Keszei, A. F. A. et al. Structure of an SspH1-PKN1 Complex Reveals the Basis for Host Substrate Recognition and Mechanism of Activation for a Bacterial E3 Ubiquitin Ligase. Mol Cell Biol 34, 362–373 (2014).

32. Keszei, A. F. A. & Sicheri, F. Mechanism of catalysis, E2 recognition, and autoinhibition for the IpaH family of bacterial E3 ubiquitin ligases. Proc. Natl. Acad. Sci. 114, 1311–1316 (2017).

33. Li, P. et al. Ubiquitination and degradation of GBPs by a Shigella effector to suppress host defence. Nature 551, 378–383 (2017).

34. Wandel, M. P. et al. GBPs Inhibit Motility of Shigella flexneri but Are Targeted for Degradation by the Bacterial Ubiquitin Ligase IpaH9.8. Cell Host Microbe 22, 507–518.e5 (2017).

35. Ji, C. et al. Structural mechanism for guanylate-binding proteins (GBPs) targeting by the Shigella E3 ligase IpaH9.8. PLoS Pathog. 15, e1007876 (2019).

36. Luchetti, G. et al. Shigella ubiquitin ligase IpaH7.8 targets gasdermin D for degradation to prevent pyroptosis and enable infection. Cell Host Microbe 29, 1521–1530.e10 (2021).

37. Wang, C. et al. Structural basis for GSDMB pore formation and its targeting by IpaH7.8. Nature 616, 590–597 (2023).

38. Hansen, J. M. et al. Pathogenic ubiquitination of GSDMB inhibits NK cell bactericidal functions. Cell 184, 3178–3191.e18 (2021).

39. Ashida, H. et al. A bacterial E3 ubiquitin ligase IpaH9.8 targets NEMO/IKKγ to dampen the host NF-κB-mediated inflammatory response. Nat Cell Biol 12, 66–73 (2010).

40. Jong, M. F. de, Liu, Z., Chen, D. & Alto, N. M. Shigella flexneri suppresses NF-κB activation by inhibiting linear ubiquitin chain ligation. Nat Microbiol 1, 16084 (2016).

41. Liu, J., et al. Mechanistic insights into the subversion of the linear ubiquitin chain assembly complex by the E3 ligase IpaH1.4 of Shigella flexneri. Proc. Natl. Acad. Sci. 119, e2116776119 (2022).

42. Kane, C. D., Schuch, R., Day, W. A. & Maurelli, A. T. MxiE Regulates Intracellular Expression of Factors Secreted by the Shigella flexneri 2a Type III Secretion System. J Bacteriol 184, 4409–4419 (2002).

43. Newton, H. J. et al. The Type III Effectors NleE and NleB from Enteropathogenic E. coli and OspZ from Shigella Block Nuclear Translocation of NF-κB p65. PLoS Pathog. 6, e1000898 (2010).

44. Zhang, L. et al. Cysteine methylation disrupts ubiquitin-chain sensing in NF-κB activation. Nature 481, 204–208 (2012).

45. Sanada, T. et al. The Shigella flexneri effector OspI deamidates UBC13 to dampen the inflammatory response. Nature 483, 623–626 (2012).

46. Ahel, J. et al. Moyamoya disease factor RNF213 is a giant E3 ligase with a dynein-like core and a distinct ubiquitin-transfer mechanism. Elife 9, e56185 (2020).

47. Szczesna, M. et al. Bacterial esterases reverse lipopolysaccharide ubiquitylation to block host immunity. Cell Host Microbe 32, 913–924.e7 (2024).

48. Zhu, K. et al. DELTEX E3 ligases ubiquitylate ADP-ribosyl modification on nucleic acids. Nucleic Acids Res. 52, 801–815 (2023).

49. Dearlove, E. L. et al. DTX3L ubiquitin ligase ubiquitinates single-stranded nucleic acids. bioRxiv 2024.04.02.587769 (2024) doi:10.1101/2024.04.02.587769.

50. Zhu, K. et al. DELTEX E3 ligases ubiquitylate ADP-ribosyl modification on protein substrates. Sci. Adv. 8, eadd4253 (2022).

51. Zou, W. & Zhang, D.-E. The Interferon-inducible Ubiquitin-protein Isopeptide Ligase (E3) EFP Also Functions as an ISG15 E3 Ligase*. J. Biol. Chem. 281, 3989–3994 (2006).

52. Kobayashi, T. et al. The Shigella OspC3 Effector Inhibits Caspase-4, Antagonizes Inflammatory Cell Death, and Promotes Epithelial Infection. Cell Host Microbe 13, 570–583 (2013).

53. Pruneda, J. N. et al. Structure of an E3:E2∼Ub Complex Reveals an Allosteric Mechanism Shared among RING/U-box Ligases. Mol. Cell 47, 933–942 (2012).

54. Guey, S. et al. Rare RNF213 variants in the C-terminal region encompassing the RING-finger domain are associated with moyamoya angiopathy in Caucasians. Eur. J. Hum. Genet. 25, 995– 1003 (2017).

55. Pinard, A. et al. Association of De Novo RNF213 Variants With Childhood Onset Moyamoya Disease and Diffuse Occlusive Vasculopathy. Neurology 96, e1783–e1791 (2021).

56. Randow, F. & Sale, J. E. Retroviral transduction of DT40. Sub-cellular biochemistry 40, 383–386 (2006).

57. Sidik, S. et al. A Shigella flexneri Virulence Plasmid Encoded Factor Controls Production of Outer Membrane Vesicles. G3 Amp 58 Genes Genomes Genetics 4, 2493–2503 (2014).

58. Schindelin, J., et al. Fiji: an open-source platform for biological-image analysis. Nat. Methods 9, 676–682 (2012).

59. Schulte, M., Olschewski, K. & Hensel, M. The protected physiological state of intracellular Salmonella enterica persisters reduces host cell-imposed stress. Commun. Biol. 4, 520 (2021).

60. Bellare, J. R., Davis, H. T., Scriven, L. E. & Talmon, Y. Controlled environment vitrification system: An improved sample preparation technique. J. Electron Microsc. Tech. 10, 87–111 (1988).

61. Russo, C. J., Scotcher, S. & Kyte, M. A precision cryostat design for manual and semi-automated cryo-plunge instruments. Rev. Sci. Instrum. 87, 114302 (2016).

62. Kimanius, D., Dong, L., Sharov, G., Nakane, T. & Scheres, S. H. W. New tools for automated cryo-EM single-particle analysis in RELION-4.0. Biochem. J. 478, 4169–4185 (2021).

63. Punjani, A., Rubinstein, J. L., Fleet, D. J. & Brubaker, M. A. cryoSPARC: algorithms for rapid unsupervised cryo-EM structure determination. Nat. Methods 14, 290–296 (2017).

64. Rohou, A. & Grigorieff, N. CTFFIND4: Fast and accurate defocus estimation from electron micrographs. J. Struct. Biol. 192, 216–221 (2015).

65. Zivanov, J., Nakane, T. & Scheres, S. H. W. Estimation of high-order aberrations and anisotropic magnification from cryo-EM data sets in RELION-3.1. IUCrJ 7, 253–267 (2020).

66. Zivanov, J., Nakane, T. & Scheres, S. H. W. A Bayesian approach to beam-induced motion correction in cryo-EM single-particle analysis. IUCrJ 6, 5–17 (2019).

67. Kimanius, D. et al. Data-driven regularisation lowers the size barrier of cryo-EM structure determination. bioRxiv 2023.10.23.563586 (2023) doi:10.1101/2023.10.23.563586.

68. Meng, E. C. et al. UCSF ChimeraX: Tools for structure building and analysis. Protein Sci. 32, e4792 (2023).

69. Hiragi, K. et al. Structural insight into the recognition of the linear ubiquitin assembly complex by Shigella E3 ligase IpaH1.4/2.5. J. Biochem. 173, 317–326 (2023).

70. Croll, T. I. ISOLDE: a physically realistic environment for model building into low-resolution electron-density maps. Acta Crystallogr. Sect. D: Struct. Biol. 74, 519–530 (2018).

71. Afonine, P. V. et al. Real-space refinement in PHENIX for cryo-EM and crystallography. Acta Crystallogr. Sect. D 74, 531–544 (2018).

72. Takeyama, K. et al. The BAL-binding Protein BBAP and Related Deltex Family Members Exhibit Ubiquitin-Protein Isopeptide Ligase Activity*. J. Biol. Chem. 278, 21930–21937 (2003).

73. Pomerantz, J. L. & Baltimore, D. NF-kappa B activation by a signaling complex containing TRAF2, TANK and TBK1, a novel IKK-related kinase. Embo J 18, 6694–6704 (1999).

74. Krissinel, E. & Henrick, K. Inference of Macromolecular Assemblies from Crystalline State. J. Mol. Biol. 372, 774–797 (2007).

75. Naydenova, K. & Russo, C. J. Measuring the effects of particle orientation to improve the efficiency of electron cryomicroscopy. Nat. Commun. 8, 629 (2017).

